# YAP/TAZ create a physical niche for the maintenance of adult neural stem cell quiescence

**DOI:** 10.1101/2025.06.27.661955

**Authors:** Laura Blasco-Chamarro, Pau García-Bolufer, Pere Duart-Abadia, Antonio Jordán-Pla, Mª Salomé Sirerol-Piquer, Pau Carrillo-Barberà, Chiara Sgattoni, Isabel Mateos-White, Cristina Gil-Sanz, Jose Manuel Morante-Redolat, Isabel Fariñas

## Abstract

Adult stem cells inhabit specialized niches where local and systemic cues regulate their behavior. In the mouse ventricular-subventricular zone (V-SVZ), neural stem cells (NSCs) dynamically transition between quiescence and activation and reside amidst unique deposits of extracellular matrix (ECM) known as fractones. We show that NSCs that enter quiescence in response to BMP4 secrete a complex ECM that, on its own, is capable of inducing NSC quiescence. This specific ECM triggers the nuclear translocation of Yes-associated protein (YAP), to induce further ECM remodeling and adhesion. Together, the BMP-ECM-YAP pathway creates a two-step mechanism where a soluble and transient quiescence-inducing signal leads to the formation of a physical niche to maintain the quiescent state. In the intact niche, YAP and its paralog TAZ (Transcriptional coactivator with PDZ-binding motif) essentially sustain quiescence by preserving fractones and the characteristic structural organization. Moreover, our findings reveal a previously unrecognized role for YAP/TAZ in quiescence.

## Introduction

Balanced quiescence and activation in adult stem cell reservoirs appear essential for preserving their long-term regenerative potential^1^. Furthermore, transcriptomic analyses have revealed that quiescence is not a passive default but a highly-regulated cellular state^2–7^. Although quiescence must be actively induced and stably maintained until activating signals prompt re-entry into the cell cycle, the molecular mechanisms specifically responsible for sustaining this state remain incompletely understood. The ability of stem cells to generate and respond to self-sustaining niche signals may, therefore, represent a critical mechanism to ensure long-term maintenance and avoid premature depletion. In the adult rodent brain, the ventricular-subventricular zone (V-SVZ) constitutes the largest and most active germinal niche, where neural stem cells (NSCs), also known as B cells, reversibly transit between quiescence and activation^3–6^. While 20% of NSCs undergoes self-renewing divisions when activated, the majority follow a consuming trajectory, producing fast-amplifying neural progenitor cells (NPCs) that commit, after a few rounds of divisions, to proliferative neuroblasts (NB1) that will differentiate into postmitotic neuroblasts (NB2). These late-stage NBs migrate to the olfactory bulb, where they terminally differentiate into interneurons that participate in olfactory discrimination^8–10^. A number of extrinsic signals have been identified as inducers of NSC quiescence^9,11–13^, but many of these cues are transient or context-dependent, raising the question of how stem cells preserve quiescence over extended periods.

Molecules mediating cell-cell and cell–extracellular matrix (ECM) interactions play a role in shaping the V-SVZ niche and have been associated with distinct activation states^14–19^. The ECM is increasingly recognized as an active regulator of stem cell behavior, acting not merely as structural support but as a dynamic and multivalent signal integrator that shapes cellular responses^20,21^. Cell–ECM interactions allow biological systems to interpret stimuli contextually, as cells exhibit different responses to the same mechanical stimulus depending on their micromechanical and biochemical surroundings^22^. Through these interactions, the ECM influences key decisions related to proliferation, differentiation, and migration across developmental and adult contexts^23^. In the V-SVZ, proteomic and ultrastructural analyses have uncovered a specialized and stiffer niche-specific ECM^24^ enriched in unique, morphologically visible extravascular basement membrane (BM) structures known as "fractones"^25–28^, which had only been previously observed in certain tumors^29^. These speckled ECM deposits, from now on “speckles”, lack an organized lamina and share key components with vascular BM, such as laminins (LMα5, β1, β2, γ1), collagen IV and VI, perlecan, agrin, nidogen, SMOC, and netrin-4, with LMα3 uniquely enriched in speckled BM^30^. Ependymal cells and NSCs contribute to speckle composition^30,31^. Disruption of *lama5* specifically in ependymal cells induces increased NSC proliferation, underscoring speckles’ regulatory role in NSC behavior^31^. Beyond their structural role, speckles also bind soluble factors, potentially modulating their availability and influencing cellular responses^25,32–34^. Furthermore, age-dependent changes in speckle number, size, and composition further support their dynamic nature and role in lifelong NSC regulation^35,36^. Nonetheless, despite extensive characterization of ECM components in the V-SVZ^24,37–39^, a comprehensive understanding of how the ECM regulates NSC states remains incomplete, including how the niche is established, the specific contribution of NSCs to ECM production, and the mechanisms underlying ECM organization within the V-SVZ.

Mechanical cues from the ECM are transduced into intracellular signals through mechanosensitive molecules such as integrins, cadherins, and Piezo channels^40,41^. These cues converge on transcriptional activators like YAP (Yes-associated protein) and TAZ (Transcriptional coactivator with PDZ-binding motif). YAP and TAZ are best known as downstream effectors of the Hippo pathway, a conserved pathway that regulates cell growth and proliferation, maintaining tissue homeostasis and controlling organ size^42–44^. They integrate mechanical and biochemical stimuli to orchestrate cell fate decisions during embryonic development, tissue regeneration, and in tumorigenesis and have been linked to the expansion of progenitor pools and the maintenance of stem-like states across multiple organs^45–47^. In the nervous system, in particular, YAP/TAZ induce the proliferation of neural progenitors during development^48–52^ and promote NSC activation in the adult subgranular zone (SGZ) of the hippocampus, another neurogenic niche^53^. However, their potential function in sustaining NSC behavior in the V-SVZ remains unexplored.

Here, we sought to elucidate the role of the ECM in regulating NSCs, focusing on how NSCs contribute to the formation and organization of the adult V-SVZ niche. Our findings provide functional evidence that quiescent NSCs actively shape the ECM composition specific to the V-SVZ. Moreover, we identify YAP/TAZ as key master regulators mediating NSC-ECM interactions, revealing a previously unrecognized role for these factors in maintaining stem cell quiescence. Our study also reveals a previously unknown role of YAP/TAZ in actively maintaining dormancy in a physiological adult stem cell population.

## Results

### Quiescent NSCs are covered by ECM

GFAP^+^ NSCs in the V-SVZ exhibit long processes that interact with vascular BM, but also with extravascular BM structures, known as fractones, that are unique to this niche^16,27,28,30,31,54^. Using immunostaining for laminins (LMs), main components of both vascular and extravascular BMs, and expansion microscopy on whole-mount V-SVZ samples, these fractones appear as speckled, amorphous, rounded structures that are closely associated with GFAP⁺ cell bodies and processes (**Figure 1A**). Transmission electron microscopy (TEM), however, reveals the distinctive fractal-like ultrastructure of these ECM deposits, showing BM enveloping glial filament-containing cellular projections, consistent with earlier characterizations^27,28^ (**Figure 1B**). In order to evaluate physical interactions of quiescent NSCs specifically, we used *in utero* electroporation (IUE) to trace NSCs that enter quiescence at mid-gestation^55,56^. Mouse embryos at embryonic (E) day 15.5 were electroporated after injection of RFP or eGFP-coding episomal plasmids into their lateral ventricles. Since plasmids become diluted with each cell division, maintaining fluorescence into adulthood depends on cells dividing infrequently^57,58^ (**Figure S1A**). In the V-SVZ of two-month-old IUE mice, sparse ependymocytes and a few NSCs retain the fluorescent label and can be clearly recognized by morphology: while ependymocytes are typically viewed as cuboidal multiciliated cells lining the ventricle, NSC are distinguished as elongated cells with varying shapes showing projections containing numerous, thin, hair-like processes (**Figures 1C, S1B**). All the NSCs that remained fluorescent fell into the deep quiescent (q) and quiescent but primed-for-activation (p) NSC populations (together referred henceforth as ‘quiescent NSCs’) when assessed with our reported flow cytometry marker panel^59^ (**Figures S1C,D**).

**Figure 1.**
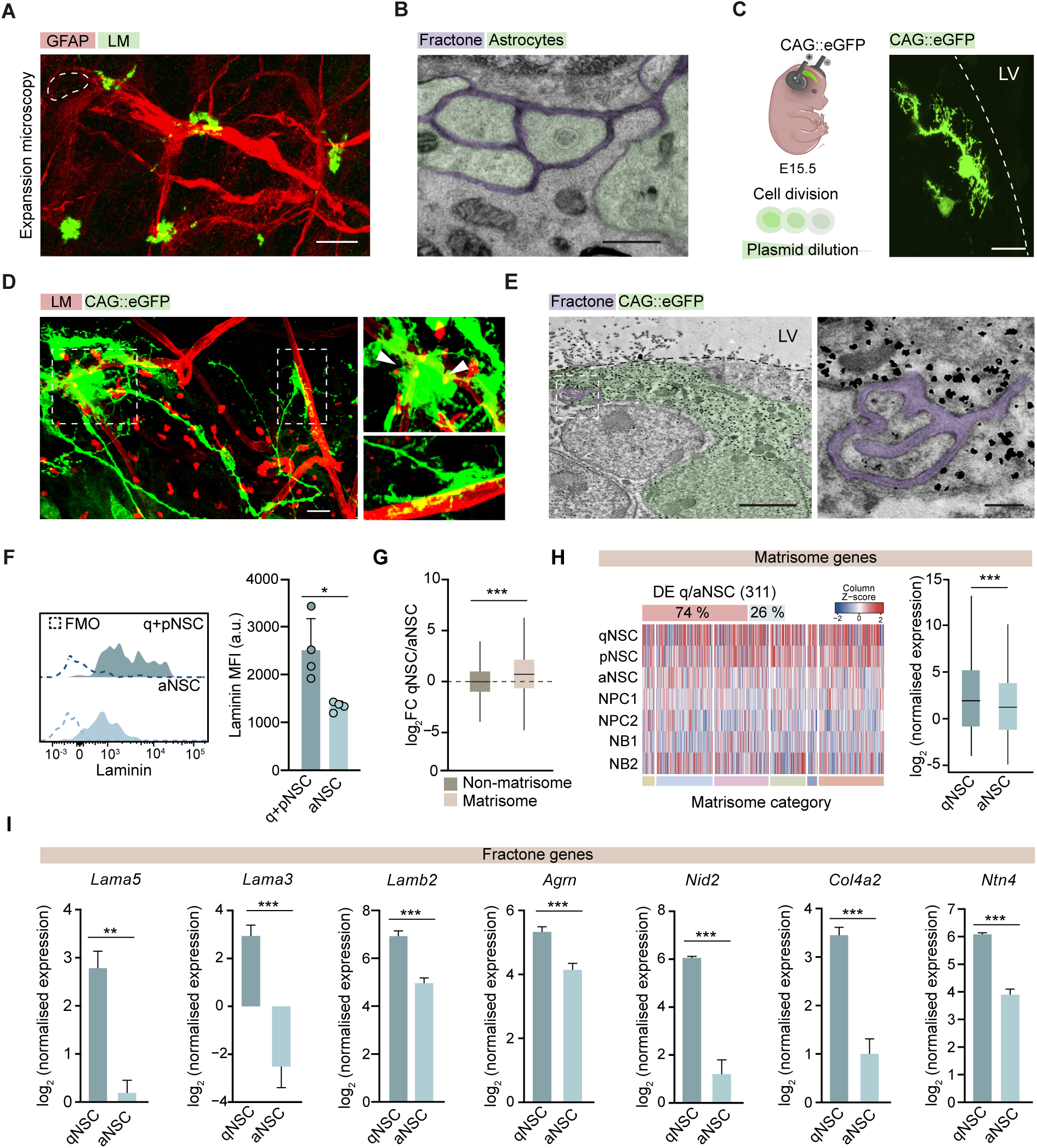
Quiescent NSCs are covered by ECM. **A.** Immunohistochemical staining of laminin (LM, green) and GFAP (red) in a whole-mount preparation of the expanded V-SVZ. The white dashed line delineates the cell nucleus. **B.** Transmission electron microscopy (TEM) images showing astrocytic projections (false-colored in green) surrounded by fractones (purple). **C.** Schematic of *in utero* electroporation with CAG::eGFP (left) and a representative confocal image of two CAG::eGFP⁺ cells in a coronal section of the V-SVZ from a 2-month-old mouse. LV: lateral ventricle. **D.** Immunohistochemical staining of optically cleared V-SVZ whole-mount preparation showing CAG::eGFP-labeled quiescent NSCs (qNSCs, green) and LM (red). Zoomed-in images show qNSC contacts with vascular and speckled basement membranes (BM). **E.** TEM images showing a qNSC (green) labeled with gold-conjugated anti-GFP antibody (black staining) in contact with fractones (purple). Right panel shows a magnified view of qNSC projections contacting fractones. **F.** Flow cytometry histograms showing LM intensity in quiescent (q and p; quiescent and primed-for-activation) and active (a) NSC populations, along with their corresponding fluorescence-minus-one (FMO) control histograms (left). LM median fluorescence intensity (MFI) in the indicated NSC populations (n=4) (right). **G.** Normalized log₂ fold change (FC) (qNSC/aNSC) in gene expression for matrisome and non-matrisome genes from bulk RNA-seq of freshly sorted NSC populations^6^. **H.** Heatmap showing the expression of matrisome genes across the neurogenic lineage, organized by matrisome categories from the Matrisome database^60^. Colors represent the different categories, in the following left-to-right order: collagens, glycoproteins, ECM regulators, ECM-affiliated proteins, proteoglycans, and secreted factors. **I.** Normalized gene expression of fractone components (*Lama5*, *Lama3*, *Lamb2*, *Agrn*, *Nid2*, *Col4a2*, *Ntn4*) in qNSCs *vs.* aNSCs from bulk RNA-seq data^6^. All graphs show mean ± SEM. ∗p < 0.05, ∗∗p < 0.01, and ∗∗∗p < 0.001. Scale bars: 20_μm (A, D); 500 nm (B); 10_μm (C); 2 μm (E, left) and 300 nm (E, right).

In optically cleared adult V-SVZ whole-mount preparations of embryonically eGFP-IUE mice, we observed that projections of quiescent NSCs terminated in LM-enriched vascular BMs and that both their cell bodies and projections were in close contact with speckled extravascular BM (**Figure 1D**). GFP immunogold staining to enable their visualization by TEM after silver enhancement, revealed very thin cytoplasmic extensions of IUE-labeled quiescent NSCs directly contacting and even embedded inside the fractal structures (**Figure 1E**). We, therefore, wondered whether the amount of extracellular LM surrounding the cells could be quantified by flow cytometry in single cell suspensions combining LM staining with our marker panel for neurogenic lineage characterization^59^. Mechanical digestion of the V-SVZ without enzymes, to prevent LM degradation, enabled us the specific detection of LM bound to the extracellular surface of the cells, with a higher intensity in quiescent NSCs (**Figure 1F**). Our results indicated a differential association of speckled BM components with the membrane of quiescent *vs*. activated NSCs.

Given the LM coverage of quiescent NSCs, we next assessed the expression of ECM molecules in these cells using the Matrisome database^60^, a meticulously curated resource integrating *in silico* and *in vivo* data, including a reported V-SVZ proteome^24^. This database provides a comprehensive, categorized set of genes encoding ECM structural components (core matrisome) and ECM-associated proteins. An analysis of our bulk RNA-seq datasets from acutely isolated V-SVZ cell fractions^6^ showed that, although qNSCs and aNSCs exhibit differences in gene expression across numerous functional categories, the fold changes (FC) in matrisome genes between these two populations were significantly greater than those observed for non-matrisome genes, suggesting distinct regulatory mechanisms for this gene subset between quiescence and activation (**Figure 1G**). The differences in gene expression between NSC states were statistically significant, with 311 differentially expressed matrisome genes, 74% of which were significantly upregulated in qNSCs (**Figure 1H**). Among them, we found multiple fractone components^30^, including LMs (*Lama5, Lama3, Lamb2*), agrin (*Agrn*), nidogen-2 (*Nid2*), collagen IV (*Col4a2*), and netrin 4 (*Ntn4*) (**Figure 1I**). As the matrisome does not include ECM receptors, we assessed the representation of *ECM-receptor interactions* in KEGG pathways comparing qNSCs *vs*. aNSCs, finding also a statistically significant enrichment in the qNSC population (p-value = 0.008). Altogether, our data suggested differential interactions of quiescent and activated NSCs with ECM and a potential role for the former in shaping the composition of extravascular BMs.

### BMP induces a quiescence proteome that includes a specific ECM-related signature

We next decided to functionally test whether quiescent NSCs indeed produce a specific ECM. Although capturing the *in vivo* qNSC state remains elusive^6^, a reversible quiescent-like state can be induced *in vitro* by BMP4. Although initially set up for hippocampal NSCs cultured on an adhesive substrate (Matrigel) and with fibroblast growth factor (FGF)^11^, the procedure has been reproduced with NSCs from the V-SVZ with similar settings^61^. However, the effect of BMPs in regular non-adherent V-SVZ cultures in the presence of FGF and epidermal growth factor (EGF) remains less well understood^62,63^. To address this, we loaded single neurosphere cells with the fluorescent division-tracer DFFDA^6^ and seeded them in uncoated plastic wells with EGF+FGF. Addition of 50 ng/ml BMP4 significantly reduced 5-ethynyl-2′-deoxyuridine (EdU) incorporation and EGFR membrane levels after 4 days *in vitro* (DIV) even under the strong mitogenic stimulation (**Figure 2A**). Consistent with the observed pro-quiescent effect, BMP4 significantly increased the proportion of DFFDA^high^ cells that retained the division-tracer (**Figure 2B**). Similar effects of BMP4 were observed when combined with either EGF or FGF alone (**Figures S2A**). Intriguingly, we found that BMP4-treated cells frequently adhered spontaneously to the uncoated plastic wells and flattened their morphology, although they remained capable of forming neurospheres upon detachment and reseeding (**Figure 2C**). Quantitative live-cell imaging revealed that BMP4 treatment triggered cell adhesion as early as 2 DIV, with a progressive increase over time (**Figure 2D, S2B; Supplementary videos 1**). Using RT-PCR, we detected elevated levels of the LM isoforms *Lama3* and *Lama2*, both of which were enriched in our RNA-seq analysis of qNSCs, in the treated cultures (**Figure 2E**). Moreover, LM secretion could be demonstrated by pan-LM immunoblotting of the culture medium (**Figure 2F**). The data suggested that adhesion induced by BMP4 might be explained by a potential NSC-mediated ECM deposition.

**Figure 2.**
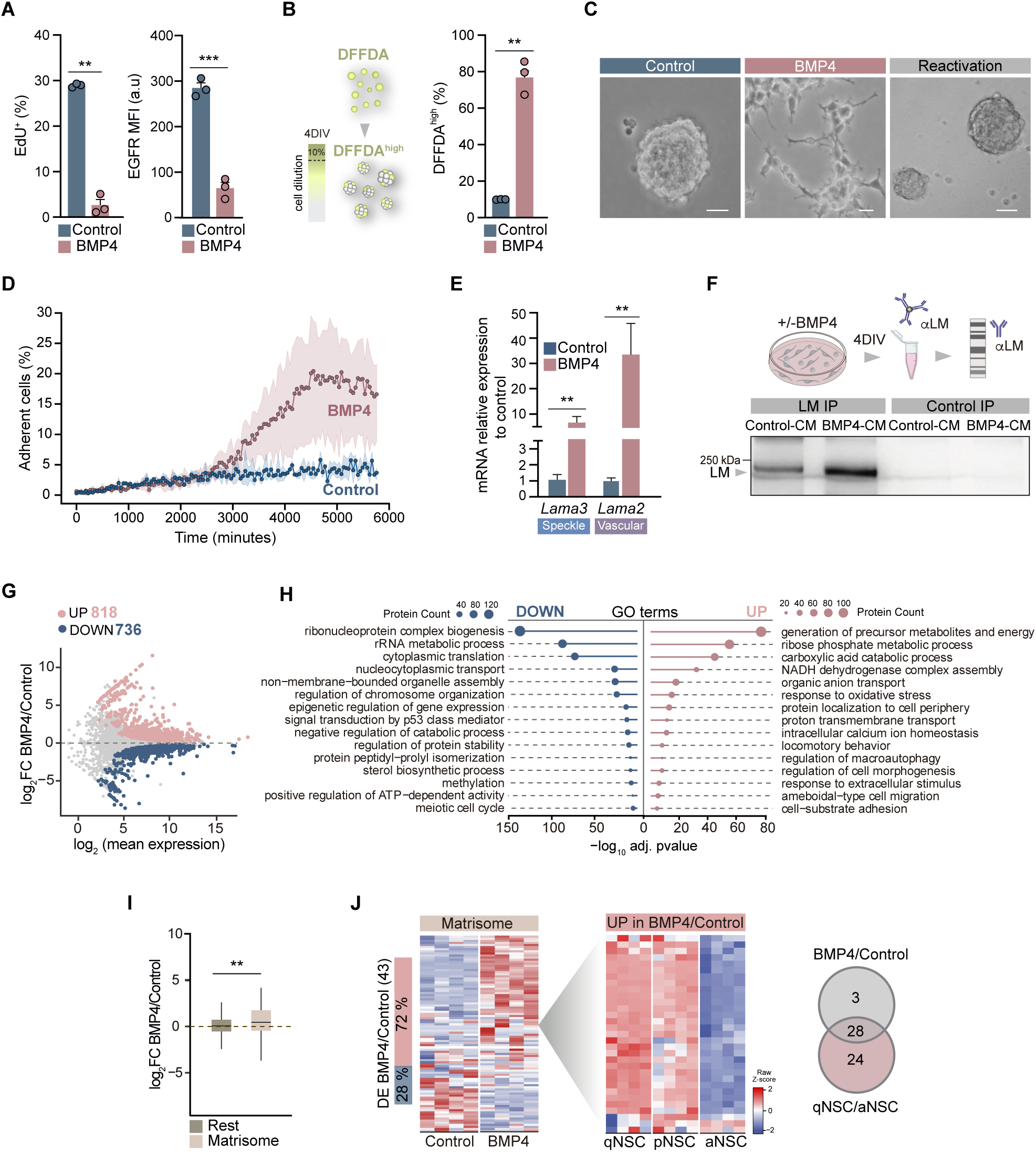
BMP induces a quiescence proteome that includes a specific ECM-related signature. **A.** Percentage of EdU⁺ cells (1_h pulse) and EGFR median fluorescence intensity (MFI) after 4_DIV with or without BMP4 (50_ng/ml) in the presence of EGF+FGF, assessed by flow cytometry (n = 3). EGFR was measured using fluorescently-labeled EGF ligands. **B.** Schematic of the strategy used to identify DFFDA^high^ cells (left). Percentage of DFFDA^high^ cells after 4_DIV with or without BMP4 (n = 3). **C.** Phase-contrast images showing NSC morphology in uncoated wells following BMP4 treatment in EGF+FGF, and after reactivation. **D.** Percentage of adherent cells over time, captured with the Incucyte® S3 live-cell imaging system over 4_DIV. Mean values and confidence intervals are shown for each time point in control and BMP4-treated NSC cultures (n = 4). **E.** Relative mRNA expression of the speckle-specific gene *Lama3* (n = 4) and the vascular-specific gene *Lama2* (n = 3) in neurosphere cultures after 4_DIV of BMP4 treatment, normalized to control samples. **F.** Immunoblot of laminin (LM) after immunoprecipitation from 4_DIV conditioned medium (CM) (combined from n = 3 cultures) using pan-LM-specific or unrelated (control) antibodies. **G.** MA plot from SWATH-MS proteomic analysis of BMP4-treated neurospheres. Unchanged proteins are shown in grey and significantly upregulated and downregulated proteins are shown in pink and blue, respectively. **H.** Top Gene Ontology (GO) Biological Process terms associated with BMP4-upregulated and - downregulated proteins. Bars represent the adjusted –log_10_ p-value of each category; circle size indicates the number of associated proteins. **I.** Normalized log_2_ fold change (FC) (BMP4/Control) in protein levels for matrisome versus non-matrisome proteins. **J.** Left: Heatmap illustrating the relative levels of matrisome proteins significantly upregulated in BMP4-treated versus control samples. Right: Heatmap displaying the *in vivo* expression of these BMP4-upregulated proteins across NSC populations. Venn diagram showing the overlap between matrisome genes significantly upregulated in qNSCs versus aNSCs and matrisome proteins significantly increased upon BMP4 treatment. All graphs show mean ± SEM. ∗p < 0.05, ∗∗p < 0.01, and ∗∗∗p < 0.001. Scale bar: 50_μm (C).

To look deeper and unbiasedly into the proteome of these BMP4-induced quiescent NSCs, we performed a quantitative differential SWATH-MS proteomic analysis of treated *vs*. untreated neurospheres. As many as 5,076 proteins were detected and principal component analysis (PCA) clearly clustered separately BMP4-treated and control samples (**Figure S2C**), with 818 and 736 proteins significantly increased and decreased, respectively (**Figure 2G; Supplementary Data 1**). To first assess whether BMP4 induced a general state resembling *in vivo* quiescence, we selected representative GO categories reported to be characteristic of qNSCs or aNSCs at the transcriptional level^2^. We then evaluated their representation in the BMP4 proteome, finding that BMP4-treated cells were significantly enriched in qNSC-associated categories and control cells in aNSC-associated ones (**Figure S2D**). Furthermore, BMP-induced proteins were found significantly upregulated at the mRNA level in acutely isolated qNSCs when compared to aNSCs *in vivo*^6^ (**Figure S2E**). The data indicated that BMP4 induces a molecular state that resembles that of qNSCs *in vivo*.

Among the top GO-terms in BMP4 upregulated proteins we found a significant enrichment in *protein localization to cell periphery, cell morphogenesis, response to extracellular stimulus or cell-substrate adhesion* (**Figure 2H**). Additionally, FCs in protein levels in BMP4 *vs*. control proteomes were significantly higher in matrisome *vs*. non-matrisome subsets (**Figure 2I**). Proteomic analysis identified 93 matrisome proteins, of which 43 were differentially expressed; notably, 72% (31 proteins) were upregulated in BMP4-treated samples compared to controls (**Figure 2J**). These included core ECM components, like proteoglycans (Bcan, Ncan) and glycoproteins (Lamb2, Creld1, Igfbp4, Sparcl1 and Vwa5a). Besides, we found multiple ECM regulators and ECM-associated proteins, including remodeling enzymes, such as Serpinh1, essential for the correct folding of procollagen, ADAM22, or cathepsins (e.g., Ctsa, Ctsb, Ctsc) (**Figure S2F; Supplementary Data 1**). Among the 31 matrisome proteins differentially increased after BMP4 treatment, 28 were differentially expressed, at the transcriptional level, in qNSCs *vs*. aNSCs *in vivo*, suggesting the existence of a specific quiescent ECM-related signature (**Figure 2J**). BMP4 treatment also upregulated receptors annotated as *ECM-receptor interactions* in KEGG pathways, such as integrin β5, that interacts with integrin αV to participate in adhesion to fibronectin and vitronectin, or in LM-binding integrin α6, reported to participate in NSC adhesion to both vascular and speckled BM *in vivo*^16,30^ (**Figure S2G**). BMP4-upregulated ECM receptors were also upregulated in qNSCs *vs*. aNSCs *in vivo* (**Figure S2G**). Our data indicated that BMP4 induces a proteome mirroring that of the quiescent state *in vivo*, as it happens in hippocampal NSCs^64^. Furthermore, this proteome includes both matrisome components and ECM receptors associated with quiescent cells, suggesting that BMP4 shapes not only quiescence, but also the ECM microenvironment and how cells interact with it.

### BMP-dependent self-generated ECM is sufficient to induce NSC quiescence through adhesion

To test for the specific effects of a quiescence-associated ECM, we let neurospheres condition their medium for 4 DIV in mitogens alone or with BMP4. Then, these conditioned media (CM), either control-CM or BMP4-CM, were supplemented with Noggin to block any residual BMP4 activity, and added to Matrigel-coated wells overnight. After thorough washing, we seeded DFFDA-loaded single neurosphere cells in normal NSC growth medium with EGF+FGF onto the coated wells (**Figure 3A**). As expected for the presence of Matrigel, cells adhered in both conditions, but only those onto BMP4-CM matrix (bMtx) stopped dividing (**Figure 3B**), suggesting that a quiescent state was being induced specifically by the matrix generated by BMP4-induced quiescent NSCs. To investigate whether adhesion was being directly induced by the bMtx, we coated untreated plastic wells overnight with BMP4-CM or control-CM with and without Noggin. Again, the wells were extensively washed the next morning and single neurosphere cells were seeded in EGF+FGF. NSCs cultured on control-CM matrix (cMtx) formed floating neurospheres, whereas those on bMtx adhered and spread across the coated surface. Besides, we found that the effect was independent of residual BMP4 activity, as similar adhesion to bMtx was observed in the presence or absence of Noggin (**Figures 3C, D; Supplementary videos 2**). To assess whether matrisome components were being deposited extracellularly and retained in cell culture wells, we used control-CM and BMP4-CM to coat plastic wells, as described above and, after washing, the coated surfaces were gently digested to perform tandem mass spectrometry (LC–MS/MS). We detected 126 matrisome proteins, 91 present in both cMtx and bMtx and 35 only detected in bMtx. Among them, we found different LM isoforms that constitute speckles (LM _5, β2, γ1) (**Figure S3**; **Supplementary Data 2**). After 4 DIV, cells on bMtx exhibited higher DFFDA tracer retention and reduced EGFR levels, quiescence features that were also independent of BMP4 (**Figure 3E**). The arrest induced by bMtx was reversible, with detached cells being able to form neurospheres (**Figures 3F**). We also investigated cycling dynamics of freshly sorted aNSCs from the V-SVZ when seeded on either cMtx or bMtx. After 24 h, a significantly higher proportion of cells remained as singlets on Noggin-blocked bMtx *vs*. cMtx, with this difference persisting at 48 and 72 h (**Figure 3G**). This experimental framework, that we have termed the induced-quiescence (iQ) assay, demonstrates that a BMP-induced specific ECM secretome is sufficient to drive a quiescent-like state in NSCs in a BMP-independent manner.

**Figure 3.**
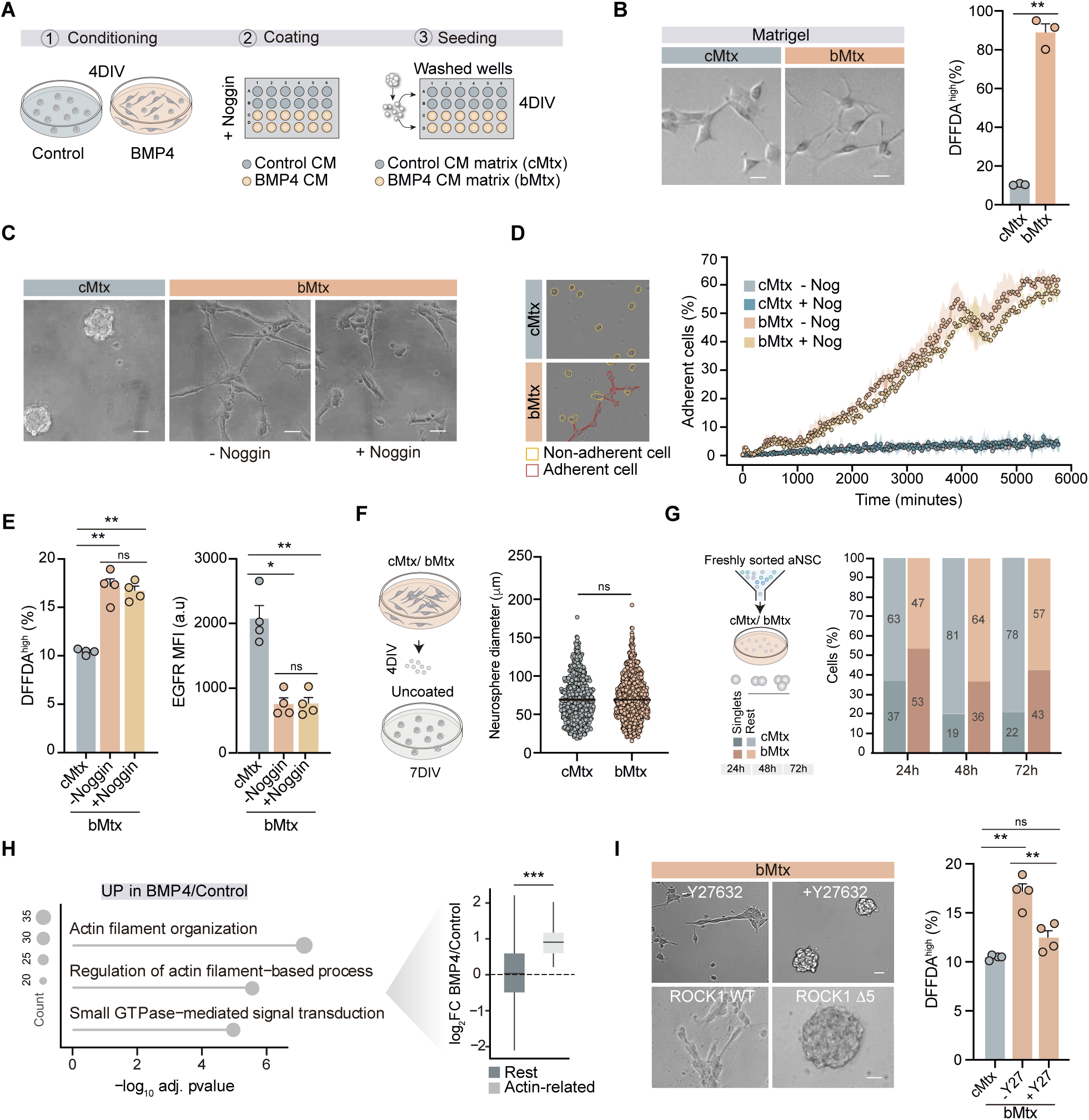
BMP-dependent self-generated ECM is sufficient to induce NSC quiescence through adhesion. **A.** Schematic overview of the experimental workflow for matrix generation and NSC culture. **B.** Left: Representative phase-contrast images of NSC cultured on Matrigel-coated wells overlayed with matrix from control-CM (cMtx) or BMP4-CM (bMtx). Right: Percentage of DFFDA^high^ cells after 4DIV in cMtx or bMtx, assessed by flow cytometry and normalized to cMtx (n = 3). **C.** Representative phase-contrast images of cells cultured on cMtx or bMtx, with or without the addition of Noggin. **D.** Left: Examples of segmented adherent and non-adherent cells. Right: Quantification of adherent cells over time, acquired using the Incucyte® S3 live-cell imaging system over 4_DIV. Mean values and confidence intervals are shown for each condition (n = 4). **E.** Percentage of DFFDA^high^ cells and EGFR median fluorescence intensity (MFI) in cMtx and bMtx, with or without Noggin (0.4 μg/ml), measured by flow cytometry. Percentages of DFFDA^high^ are normalized to cMtx (n = 4). **F.** Schematic of the experimental design (left). Neurosphere diameters measured after 7 DIV from detached cells previously cultured for 4 DIV in cMtx or bMtx (right). **G.** Freshly sorted activated NSCs (aNSCs) cultured on cMtx or bMtx. Bar graph displays the proportion of cells that remained as singlets (dark orange/blue) or underwent at least one division (light orange/blue) at 24, 48, and 72 _h. **H.** Gene ontology (GO) enrichment analysis of selected categories in BMP4 *vs*. control conditions. Left: –log_10_ adjusted *p*-value and protein counts (circle size). Right: Normalized log_2_ fold change (FC) (BMP4/control) in protein levels for proteins associated with actin-related categories *vs*. all others. **I.** Representative phase-contrast images of NSCs cultured on bMtx following treatment with the ROCK inhibitor Y27632 (25 μM) or nucleofection with a dominant-negative ROCK1 mutant (Δ5). Untreated cells and cells nucleofected with ROCK1 wild-type (WT) are also shown. Right: Quantification of DFFDA^high^ cells cultured on bMtx with or without Y-27632, normalized to cMtx (n = 4). Data for cMtx and bMtx + Noggin are replotted from panel E. All graphs show mean ± SEM. ∗p < 0.05, ∗∗p < 0.01, and ∗∗∗p < 0.001. Scale bars: 10 μm (B); 50 μm (cMtx) and 10 μm (bMtx) (D); 100_μm (top) and 20_μm (bottom) (I).

Quiescence-to-activation transitions require major changes in transcriptional programs^2^. Given the complexity of the bMtx, we focused on intracellular signaling as a more integrative readout of ECM-driven responses. Notably, actin organization and small GTPase signaling, key pathways linking ECM cues to nuclear responses, were enriched in BMP4-treated cells (**Figure 3H**). Besides, proteins associated with the actin cytoskeleton showed significantly higher FCs in the BMP *vs*. control condition compared to overall proteome changes, suggesting that BMP signaling specifically enhances actin-related processes (**Figure 3H**). The actin cytoskeleton not only provides structural support and facilitates cell movement but also plays a key role as a mechanotransducer, converting physical signals from the ECM into biochemical signals that activate signaling pathways (e.g., Rho GTPases, FAK) and transcription factors (e.g., YAP/TAZ, NF-κB)^40^. Actin stress fiber formation is regulated by the Rho GTPase family^65,66^, with Rho-associated coiled-coil containing kinase (ROCK) serving as a key downstream effector. Thus, to assess the role of the actin cytoskeleton in NSC adhesion and quiescence, we seeded single neurospheres onto bMtx in the presence or absence of the ROCK inhibitor Y27632 or after nucleofection with a dominant-negative ROCK1 mutant (Δ5). After 4 DIV, cells treated with Y27632 or transfected with ROCK Δ5 failed to adhere to bMtx in the iQ assay and, instead, formed floating neurospheres that proliferated to the extent of the cMtx condition (**Figure 3I**). These findings indicated that ROCK-dependent actin cytoskeletal dynamics are necessary for NSC adhesion to the BMP-induced matrix and the subsequent induction of quiescence.

### A pro-quiescence self-generated ECM triggers the nuclear translocation and activity of YAP

The transcriptional regulator YAP is typically involved in translating microenvironmental physical sensing to gene expression and its movement to the nuclear compartment relies on cytoskeletal contractility and ROCK activity, among other mechanisms^43,67–70^. When YAP translocates to the nucleus in its unphosphorylated form, it interacts with TEAD transcription factors to regulate gene expression^71^. To assess the potential role of this co-activator in the responses to NSC self-assembled ECM we transfected neurosphere cells with GFP-IRES-YAP1-TEAD-P-H2B-mCherry, a bicistronic construct carrying an mCherry-fused histone2B (H2B-mCherry) under the TEAD-responsive promoter and a GFP-IRES-YAP1 under a CMV promoter, designed to monitor YAP/TEAD transcriptional activity in real-time^72^. Culture of the transfected cells on cMtx, bMtx, or Matrigel as a control of adhesion, in the presence of Noggin, indicated that YAP/TEAD-dependent transcriptional activity was specifically increased on bMtx (**Figure 4A**). Notably, we found higher active YAP nuclear immunofluorescent levels in non-proliferating (Ki67^-^) *vs*. proliferating (Ki67^+^) cells adhered to Noggin-blocked bMtx (**Figures 4B**). The results indicated that a pro-quiescence ECM induces YAP nuclear activity.

**Figure 4.**
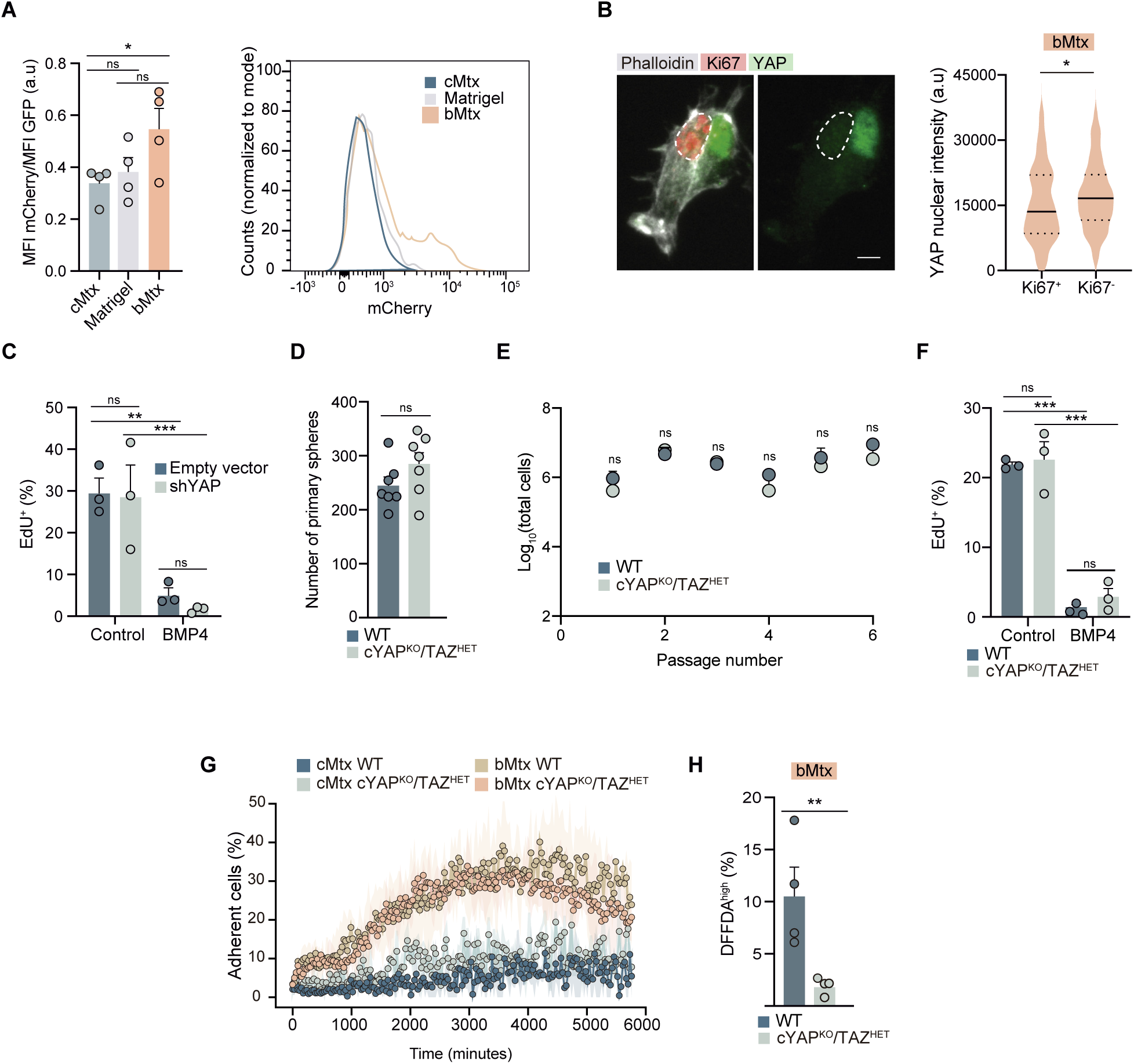
A pro-quiescence self-generated ECM triggers the nuclear translocation and activity of YAP. **A.** YAP/TEAD reporter (FLAG-YAP1-TEAD-P-H2B-mCherry) activity, assessed as mCherry mean fluorescence intensity (MFI) normalized to GFP MFI in GFP^+^ cells 24_h post-transfection, measured by flow cytometry (n = 4). A representative mCherry fluorescence histogram is shown. **B.** Left: confocal fluorescence image of NSCs cultured on pre-coated Matrigel surfaces and overlaid with bMtx for 4DIV and stained with YAP (green), Ki67 (red) and phalloidin (grey). A Ki67^+^ cell nucleus is delineated. Right: violin plot showing quantification of YAP nuclear mean fluorescence intensity in Ki67^+^ and Ki67^-^ nuclei. Median and upper and lower quartiles are indicated. **C.** Percentage of EdU^+^ cells 48 h after transfection with shYAP or empty vector, calculated among transfected (GFP^+^) cells cultured with or without BMP4 (n = 3). **D.** Number of primary neurospheres formed from WT and YAP^KO^/TAZ^HET^ cells after 10 DIV (n = 7). **E.** Growth curve of WT and cYAP^KO^/TAZ^HET^ from passage 1 to 5 (n = 3). **F.** Quantification of percentage of EdU^+^ cells (1-h pulse) in WT and cYAP^KO^/TAZ^HET^ cells, after 4 DIV with or without BMP4 (n = 3). **G.** Quantification of the percentage of adherent cells over time, acquired using the Incucyte® S3 live-cell imaging system over 4_DIV on cMtx and bMtx in WT (n = 3) and cYAP^KO^/TAZ^HET^ cells (n = 2). **H.** Quantification of the percentage of DFFDA^high^ cells in WT and cYAP^KO^/TAZ^HET^ cells after 4 DIV on bMtx (n = 4). All graphs show mean ± SEM. ∗p < 0.05, ∗∗p < 0.01, and ∗∗∗p < 0.001. Scale bar: 5 μm (B).

In order to evaluate the potential YAP link between the ECM adhesion and proliferative arrest, we first tested the role of YAP in floating neurospheres. Cells transfected with a *Yap* shRNA for 48 h incorporated EdU to the same extent as cells transfected with the empty vector, showing that YAP did not basally affect proliferation. Besides, we found that cells responded normally to the effect of BMP4 in lowering the proportion of EdU incorporation (**Figure 4C**). Although not fully redundant, YAP and transcriptional co-activator with PDZ-binding motif (TAZ) are paralog proteins with structural and functional similarities, displaying overlapping functions that can compensate for each other^73^. In line with the reported phenotypes for multiple *Yap/Taz* mutant mouse models^74^, we decided to use cells from mice without YAP and only one gene dose of TAZ. For perinatal recombination in GFAP^+^ NSCs, mutant mice were obtained by crossing mGFAP-Cre mice^75^ with floxed *Yap/Taz* mice, which possess loxP sites flanking exons 2 of both *Yap1* and *Wwtr1* (*Taz*) genes^76^. Conditional cYAP^KO^/TAZ^HET^ mice showed indistinguishable primary neurosphere number and growth rates (**Figures 4D,E**). Besides, we observed normal EdU labeling after a 1 h-pulse in floating neurospheres cultured in EGF+FGF, and BMP4 reduced EdU incorporation in the same proportion as in wild-type (WT) ones (**Figure 4F**). The data indicated that BMP does not require YAP/TAZ to induce quiescence in floating neurospheres. Then, we evaluated the role of YAP/TAZ in the context of adhesion to Noggin-blocked bMtx. We found that, while cYAP^KO^/TAZ^HET^ cells attached and spread on bMtx (**Figure 4G; Supplementary videos 3**), they could not enter quiescence as efficiently, showing a reduced proportion of DFFDA^high^ cells (**Figure 4H**). These loss-of-function experiments indicated that YAP/TAZ signaling is required to link ECM adhesion to quiescence induction.

### YAP/TAZ participate in ECM production and sustain cell-ECM adhesion

Given that bMtx induces YAP translocation to the nucleus to perform its transcriptional function, we decided to test the effect of forcing YAP activity by transfecting neurosphere cells with a constitutively active YAP (YAP-5SA), with five mutated serines that hinder its inactivation by LATS kinases^77^. YAP gain-of-function reduced proliferation in neurospheres, assessed by EdU incorporation after 48 h. This cytostatic effect was TEAD-dependent, as cells transfected with a TEAD-binding-deficient YAP construct (YAP-5SA/S94A) did not show any effect and resembled the empty vector (pBabe)-transfected controls (**Figure 5A**). The data indicated that forcing YAP activation is sufficient to induce quiescence, in contrast to reported YAP actions^44-53^. We also observed that some YAP-5SA-overexpressing cells were attaching to the uncoated plastic plates (**Figure 5B**), making us hypothesize that YAP might be regulating ECM deposition to reinforce the adhesive responses.

**Figure 5.**
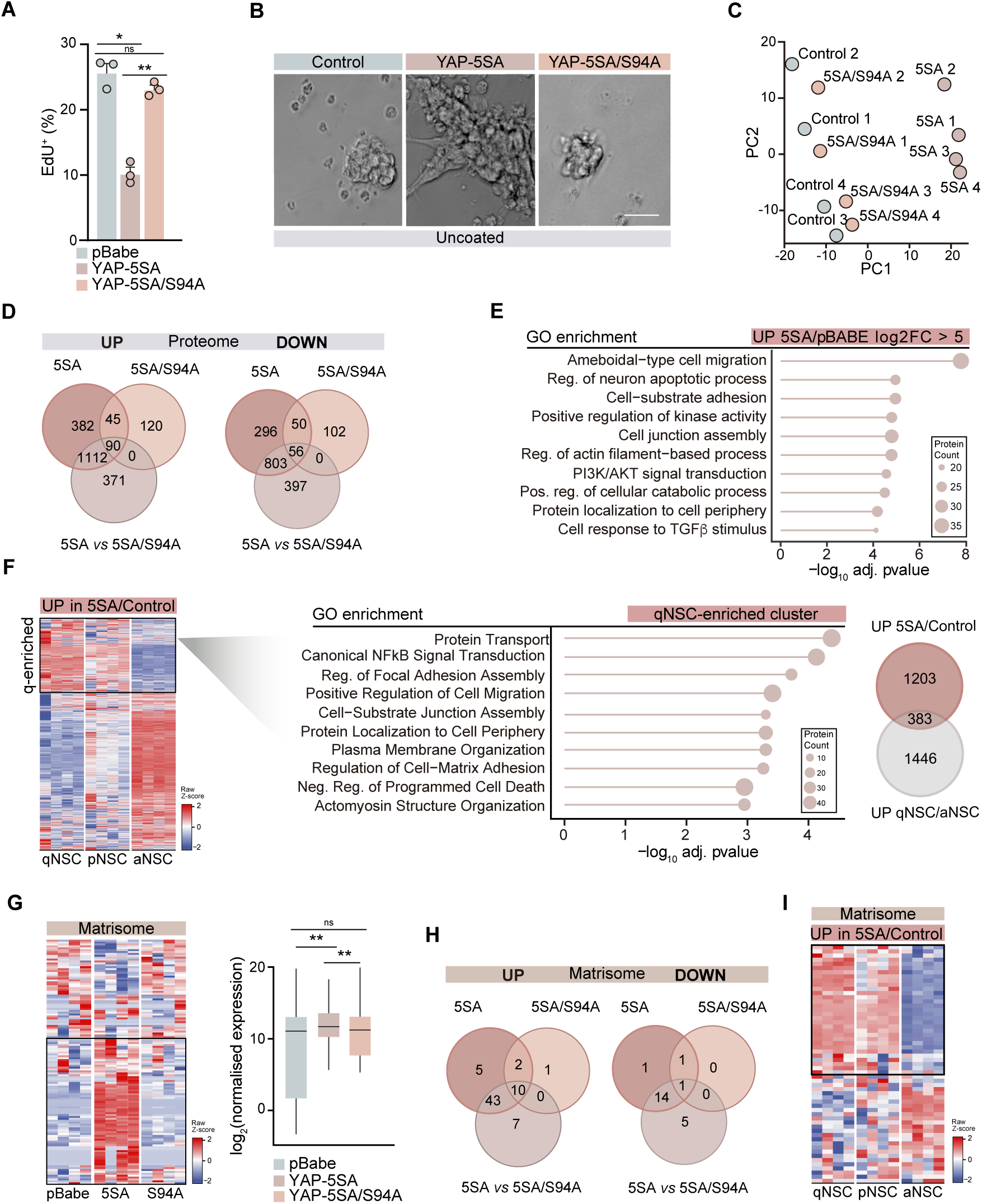
YAP/TAZ participate in ECM production and sustain cell-ECM adhesion. **A.** Quantification of percentage of EdU^+^ cells (1h-pulse), assessed by flow cytometry, in NSCs on uncoated wells, 24 h after nucleofection with constitutively active YAP (5SA), a TEAD-binding-deficient mutant (5SA/S94A), or empty vector (pBabe) (n = 3). **B.** Representative phase-contrast images of NSC nucleofected with the indicated YAP constructs and cultured on uncoated wells for 24 h. **C.** Principal component analysis (PCA) of SWATH-MS proteomic profiles from cells expressing 5SA, 5SA/S94A, or control vector (pBabe). **D.** Venn diagrams showing the overlap of significantly upregulated and downregulated proteins in 5SA and 5SA/S94A compared to pBabe, and between 5SA and 5SA/S94A. **E**. Gene ontology (GO) enrichment analysis (Biological Process) of proteins highly enriched (log₂FC > 5) in 5SA *vs.* pBabe conditions. **F.** Heatmap of proteins upregulated in 5SA *vs*. pBabe and their expression across NSC populations in bulk RNA-seq data^6^. GO enrichment analysis (Biological Process) of genes enriched in qNSCs. Venn diagram showing the overlap between proteins upregulated in 5SA/pBabe and genes enriched in qNSCs *vs*. aNSCs. **G.** Heatmap showing the expression of matrisome-associated proteins in cells expressing 5SA, 5SA/S94A, and pBabe. Right: log_2_ normalised expression of matrisome genes in pBabe, YAP-5SA and YAP-5SA/S94A. **H.** Venn diagrams showing the intersection of differentially expressed matrisome proteins among the three conditions. **I.** Heatmap showing the gene expression of 5SA/pBabe-upregulated matrisome proteins in NSC populations *in vivo*^6^. Graphs show mean ± SEM. ∗p < 0.05, ∗∗p < 0.01, and ∗∗∗p < 0.001. Scale bar: 50 μm (B).

To investigate potential YAP-dependent ECM shaping by NSCs, we electroporated single neurosphere cells with the YAP-5SA and YAP-5SA/S94A constructs together with an eGFP plasmid and, after 48 h, we sorted transfected cells for diaPASEF-MS proteomic analysis. PCA further supported the notion that YAP function in NSCs is largely TEAD-dependent, as YAP-5SA samples clustered separately from the control group, while YAP-5SA/S94A samples overlapped with controls (**Figure 5C**). The proteomic analysis identified nearly 7,000 proteins, with 1,629 and 1,573 significantly upregulated, and 1,205 and 1,256 downregulated proteins in YAP-5SA cells compared to pBabe and to YAP-5SA/S94A, respectively (**Figure 5D; Supplementary Data 3**). In contrast, the comparison between YAP-5SA/S94A and the control (pBabe) revealed only 255 upregulated and 208 downregulated proteins, further supporting that YAP activity in NSCs is predominantly mediated through TEAD-dependent mechanisms (**Figure 5D**). GO analysis revealed that *cell-substrate adhesion, cell junction assembly* or *regulation of actin filament-based process* were among the biological processes most affected by YAP gain-of-function (5SA) (FC>5) (**Figure 5E**). However, differently from the BMP4 proteome, which globally overlapped with qNSC *in vivo*, YAP gain-of-function proteomics only partially recapitulated qNSC signatures, with 32% of the increased proteins in YAP-5SA/pBabe being differentially upregulated at the mRNA level between qNSCs and aNSCs (**Figure 5F**). When evaluating GO analysis of this specific cluster of genes, we found that GOs related to *focal adhesion assembly, regulation of cell substrate junction assembly* or *actomyosin structure organization* were among the most enriched categories. The data together indicated that YAP modulates a specific transcriptional signature in qNSCs that is mostly related to cell-ECM interactions, in line with our *in vitro* data.

A specific focus on ECM proteins showed an enrichment in proteins from different matrisome categories when YAP was constitutively active in NSCs (**Figure 5G**). Specifically, differential expression analyses identified 129 ECM-related proteins, with 60 increased and 17 decreased in YAP-5SA compared to controls (**Figure 5H, Supplementary Data 3**). Among the ECM proteins regulated by YAP, we found increased BM components, including fibril-forming collagens (e.g., V and XVIII), which are implicated in tissue stiffness^78^, fibronectin or fibrillin, and LM isoforms β1 and γ1 (**Figure S4A**). Multiple ECM remodelers were also regulated by YAP, including tissue inhibitor of metalloproteinases (Timp2 and Timp3), that inhibit ECM degradation, and proteins associated with collagen cross-linking and folding (Loxl2, Plod1/2/3, P4ha1, and Serpinh1) (**Figure S4A**). To study whether YAP could also regulate ECM deposition *in vivo*, we assessed mRNA expression of YAP-regulated matrisome proteins in quiescent and activated NSCs, finding that 42% of the matrisome proteins increased in YAP-5SA/pBabe overexpressing cells were also increased, at the mRNA level, in qNSCs compared to aNSCs (**Figure 5I, S4B**). We also found different ECM receptors upregulated, including focal-adhesion mediators integrin α1, α3 and α5, β1, LM and collagen-binding CD44 or dystroglycan 1 (Dag1) that were also upregulated, at the transcriptional level, in qNSC *vs*. aNSCs (**Figure S4C**). Besides integrins, we found that constitutive activation of YAP significantly increased focal adhesion regulators (e.g. vinculin, talin, zyxin, and paxillin) some of which were also found commonly upregulated in qNSCs (**Figure S4D**). Our findings demonstrate that YAP contributes to ECM remodeling and modulates ECM-dependent signaling in NSCs *in vitro*, while orchestrating a specific adhesion-associated transcriptional program in qNSCs *in vivo*.

Integrating both BMP4 and YAP gain-of-function proteomes, we found that matrisome proteins significantly increased in each case were almost mutually exclusive (**Figures S4E**). Only a few commonly regulated proteins were identified, including Annexins and S100 families, which have been reported as characteristic of the V-SVZ proteome when compared to the cortex^24^. Our data together suggests a two-step process in the induction of quiescence: (1) BMP4 induces quiescence and drives the deposition of an ECM that favors YAP activation, and (2) YAP further shapes ECM deposition and reinforces cell interaction with the ECM.

### YAP/TAZ regulate the niche ECM structure to preserve NSC quiescence

As a first approach to assess whether YAP-regulated transcription is active in the *in vivo* V-SVZ niche, we decided to take advantage of a published YAP-conserved transcriptional signature^79^. While this signature has been shown to be enriched in active NSCs in the subgranular zone niche^53^, we found that 60% of the YAP-regulated genes were found overexpressed in qNSCs in the V-SVZ (**Figure S5A**). This was confirmed using a YAP chromatin immunoprecipitation sequencing (ChIP-seq) dataset^69^, which also showed a cluster of YAP-regulated genes upregulated in qNSCs compared to aNSCs (**Figure S5B**). In this cluster, *cell substrate adhesion*, *small GTPase signal transduction* or *cell morphogenesis* were again among the most represented GO categories (**Figure S5C**). In line with these data, we found that non-proliferative (Ki67⁻) GFAP^+^ cells in the intact niche were enriched in nuclear unphosphorylated (active) YAP compared to proliferative (Ki67^+^) cells (**Figure 6A, S5D**).

**Figure 6.**
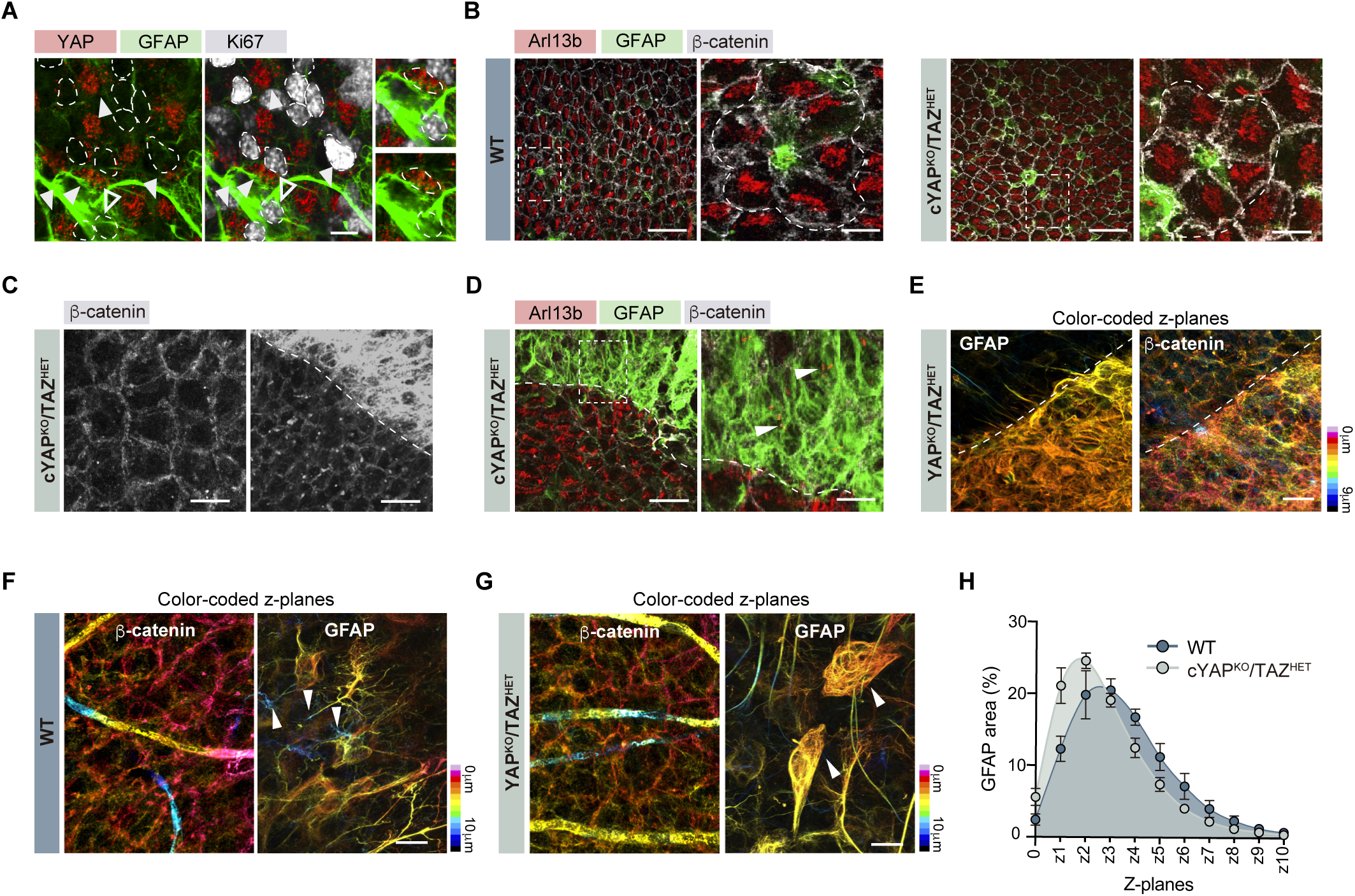
YAP/TAZ regulate niche structure and NSC apico-basal distribution. **A.** Representative confocal images of a V-SVZ whole-mount immunostained for active YAP (red), GFAP (green), and Ki67 (grey). White dashed line outline Ki67⁺ nuclei. Empty arrowheads indicate GFAP⁺/Ki67⁺ cells; filled (white) arrowheads indicate GFAP⁺/Ki67⁻ cells. **B.** Representative confocal images of immunostaining of cilium marker Arl13b (red), GFAP (green) and β-catenin (grey) in WT and cYAP^KO^/TAZ^HET^ mice. Right panels display magnified views of the areas outlined with a white dashed line. Pinwheel-like structures are delineated. **C.** Representative images of β-catenin (grey) immunostaining in cYAP^KO^/TAZ^HET^ whole-mount, showing regions with preserved β-catenin organization (left) and areas with disorganized β-catenin architecture (right). The white dashed line marks the boundary between normal and structurally altered regions. **D.** Representative confocal image of a structurally altered region showing GFAP^+^ cells with Arl13b^+^ primary cilia. Right panel shows a higher magnification of the outlined area; white arrowheads highlight Arl13b^+^ cilia. Normal and disorganized regions are delimited by a white dashed line. **E.** High-resolution color-coded z-projection images depicting spatial distribution of GFAP (left) and β-catenin (right) immunostainings. White dashed line separates normal and structurally altered areas. Distance from the ependymal surface (0_μm) is represented with a color-coded scale. **F.** Z-projection color-coded images of β-catenin (left) and GFAP (right) in WT preparations. White arrowheads indicate NSC processes projecting toward basal planes. Color scales represent distance from the ependymal cell layer (0_μm). **G.** Equivalent z-projection color-coded images in cYAP^KO^/TAZ^HET^ samples. White arrowheads indicate enlarged GFAP⁺ cells occupying the ependymal plane. **H.** Quantification of the percentage of GFAP⁺ area *per* z-plane normalized to total GFAP area in WT (n=3) and cYAP^KO^/TAZ^HET^ samples (n = 4), using the ependymal surface as the reference plane (0_μm). Interaction between genotype and z-plane was statistically significant (p < 0.0001). Scale bars: 5 μm (A); 50 μm (right) and 10 μm (left) (B); 10 μm (left) and 20 μm (right) (C); 20 μm (right) and 10 μm (left) (D); 10 μm (E, F, G).

To evaluate the *in vivo* role of YAP, we analyzed whole-mount preparations of the V-SVZ of 2-month-old cYAP^KO^/TAZ^HET^ and WT mice. In WT preparations, GFAP^+^ B1 NSCs were aligned with ependymal cells, forming the "pinwheel" pattern characteristic of this niche^80^. This organization becomes apparent when immunostained for β-catenin, which highlights the membrane adhesive complexes between the involved cells and for cilia components, which distinctly differentiate the multiciliated surface of the ependymal cells from the short primary cilia of B1 NSCs that co-localize with GFAP in the middle of the pinwheel^80^ (**Figure 6B**). Most of the V-SVZ in cYAP^KO^/TAZ^HET^ mutants displayed the characteristic β-catenin^+^ tiled-like structures, indicating that the ependyma was normally organized (**Figure 6B**). However, this organization appeared disrupted in certain regions of the cYAP^KO^/TAZ^HET^ mice (**Figure 6C**). In those “disorganized” zones, ependymal cells were substituted by supernumerary densely packed GFAP^+^ cells with Arl13b^+^ primary cilia (**Figure 6D**). Notably, these cells exhibited a loss of their typical bipolar morphology, as they were located within the same plane as the ependymal cells, but were still positive for β-catenin, suggesting the preservation of adherens junctions between cells (**Figure 6E)**. Using TEM analysis, we also observed regions with an accumulation of B cells forming cell-cell junctions among them and with nearby ependymal cells, exhibiting a high density of glial filaments and multiple interwoven and overlapping projections, as also observed by immunostaining. These regions exhibited increased cellularity (more B cells and progenitors) and a higher number of cells in prophase, supporting the notion that these areas harbor highly proliferative NSCs at the ependymal plane (**Figure S6A-D**).

Considering that these regions comprised only a small portion of the whole V-SVZ area (3.18 ± 2.10%), we questioned whether these visibly altered zones might simply be more advanced manifestations of a progressive process already underway across the V-SVZ tissue. Indeed, in regions where the pinwheel structure appeared intact, we also found abnormally shaped GFAP^+^ cells with enlarged cell bodies, abundant intermediate filaments, and thickened projections throughout the whole V-SVZ (**Figure 6F,G**). This greatly contrasted with the more uniform and consistent morphology of WT cells, which were typically bipolar with smaller cell bodies and narrower, less prominent basal projections (**Figure 6F,G**). Besides, mutant hypertrophic GFAP^+^ cells were also found to lose their normal apico-basal orientation and to localize closer to and even within the ependymal plane, as confirmed by measurements in each z-plane using high-resolution confocal microscopy with color-coded images (**Figure 6G, H**). TEM images confirmed this misorientation and loss of polarity, including the misalignment of the primary cilium. Ultrastructural analyses revealed complete disorganization in specific regions of the niche, with a loss of cell polarity (**Figure S6E-J**), finding cells dividing towards the ventricle and even within the ventricle (**Figure S6K**). This abnormal positioning, observed along the V-SVZ in patches of varying sizes, suggested a progressive phenotype, ranging from small clusters of cells that lose their normal location to larger groups that colonize the ependymal layer.

Based on our previous data, we hypothesized that this phenomenon might be mediated by a disrupted ECM regulated by YAP/TAZ. Given that our previous proteomic analysis showed YAP regulation of speckled BM components in a gain-of-function context and considering the importance of these structures in NSC regulation, we aimed to assess the presence of speckles and NSC interactions with speckles across the different regions identified. We immunostained cYAP^KO^/TAZ^HET^ mice V-SVZ whole-mounts for LM and first quantified speckle number and size. For that, we distinguished between the smaller more extreme regions of GFAP patches with altered β-catenin ependyma structure and the rest of the V-SVZ that showed apparently normal β-catenin ependymal tiled-like structures. We found that most regions of the V-SVZ still contained speckles; however, their number and size were reduced compared to WT mice, suggesting a role for YAP in the normal formation of speckles within the V-SVZ (**Figure 7A, B**). Notably, regions exhibiting GFAP patches were completely devoid of speckles (**Figure 7C**). This absence was also confirmed at the ultrastructural level, as no fractal structures were observed in TEM images of these areas (**Figure 7D**).

**Figure 7.**
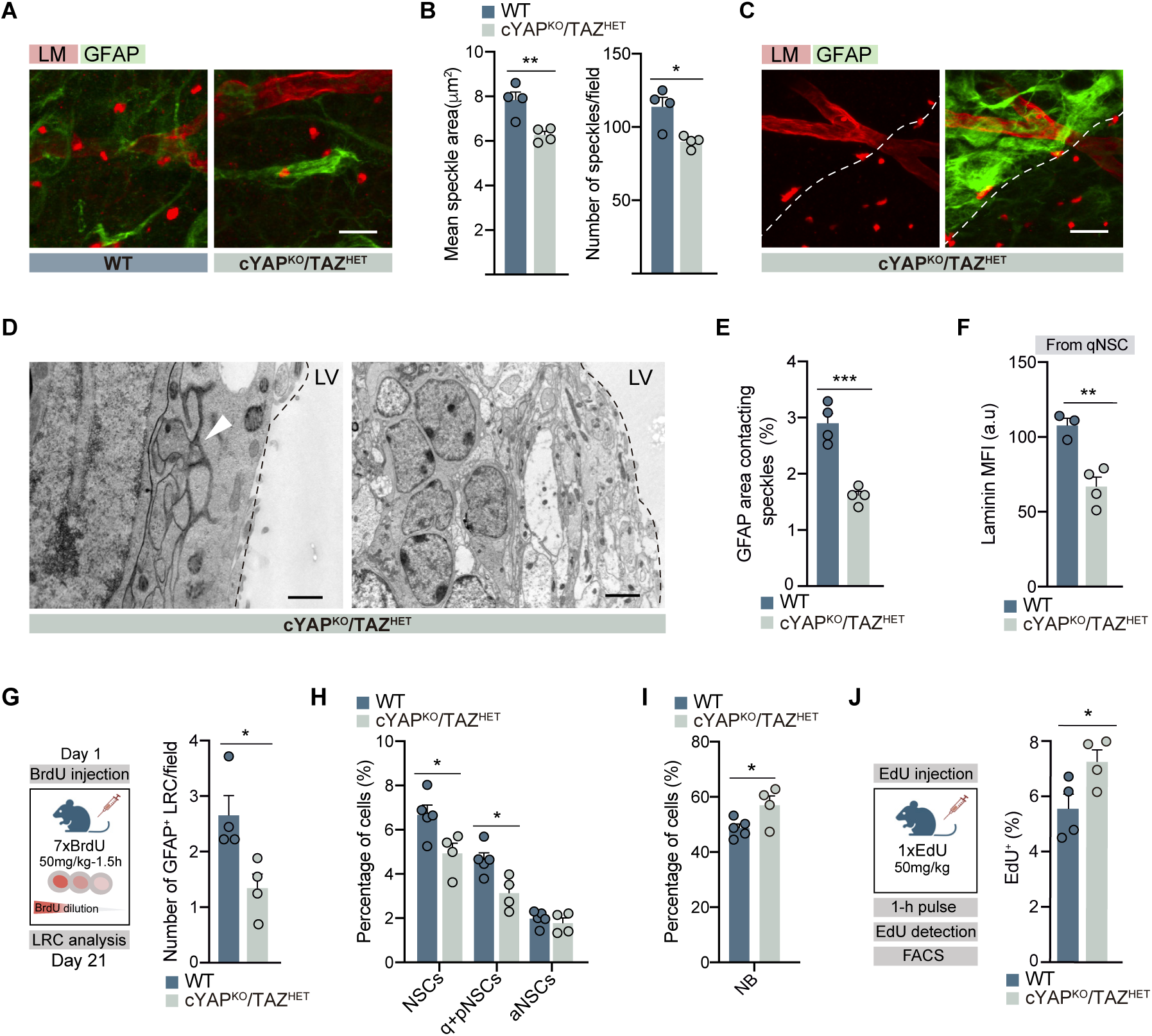
YAP/TAZ contribute to speckle formation and the maintenance of NSC quiescence. **A.** Representative confocal images of LM and GFAP immunostaining in V-SVZ whole-mounts of WT and cYAP^KO^/TAZ^HET^ mice, in regions with preserved β-catenin structures. **B.** Quantification of mean speckle area and speckle number per field (1,35 x 10^-4^ mm^3^) in normally structured areas of WT and cYAP^KO^/TAZ^HET^. **C.** Representative confocal images of LM and GFAP in V-SVZ whole-mount in structurally altered regions of WT and cYAP^KO^/TAZ^HET^. Altered regions are located on the left-hand side of the white dashed line. **D.** Transmission electron microscopy (TEM) images of normal (left) and structurally altered (right) regions in cYAP^KO^/TAZ^HET^. White arrowhead indicates fractal-like ECM structures in normally organized regions. LV: lateral ventricle. **E.** Quantification of GFAP⁺ area in contact with LM⁺ speckles, normalized to GFAP total area, in normally structured regions of WT and cYAP^KO^/TAZ^HET^. **F.** Median fluorescence intensity (MFI) of LM in quiescent NSCs (qNSCs), assessed by flow cytometry in WT and cYAP^KO^/TAZ^HET^ mice. **G.** Schematic overview of the experimental pipeline. Quantification of GFAP⁺ label-retaining cells (LRC) per field (7,2 x 10^-4^ mm^3^) in WT and cYAP^KO^/TAZ^HET^ mice. **H.** Percentage of total NSCs, quiescent (q) and primed-for-activation (p) NSCs and active (a) NSCs in the V-SVZ of WT and cYAP^KO^/TAZ^HET^ mice assessed by flow cytometry. **I.** Flow cytometry quantification of total neuroblast (NB) population in the V-SVZ of WT and cYAP^KO^/TAZ^HET^ mice. **J.** Scheme of the experimental setup (left) and flow cytometry quantification of EdU^+^ cells (1-h pulse) in the neurogenic lineage in WT and cYAP^KO^/TAZ^HET^ mice. All graphs show mean ± SEM. ∗p < 0.05, ∗∗p < 0.01, and ∗∗∗p < 0.001. Scale bars: 10 μm (A, C); 500 nm (right) and 2 μm (left) (D).

Then, we set out to quantify NSC-speckle contacts. We found that cYAP^KO^/TAZ^HET^ mice showed reduced speckle contact areas, quantified as the percentage of GFAP⁺ area in contact with LM⁺ speckles, normalized to total GFAP area (**Figure 7E**). We then assessed LM association with qNSCs by flow cytometry and, again, found reduced extracellular LM in cYAP^KO^/TAZ^HET^ qNSCs (**Figure 7F**). To examine the functional consequences of this disturbed ECM in NSC regulation, we injected mice with BrdU (50 mg/kg, 7 doses every 1.5 h) and analyzed label retaining cells (LRC), cells with slow-cycling dynamics that still retain the analogs 21 days post-injection. With this approach, we found a significant reduction in the number of GFAP⁺ BrdU⁺ LRCs in cYAP^KO^/TAZ^HET^ mice whole-mounts compared to control mice, suggesting reduced number of slow-cycling cells (**Figure 7G**). In line with this, flow cytometry assessment of the neurogenic lineage revealed a reduction in the NSC pool, which was specifically attributed to a decline in the quiescent NSC population (**Figure 7H**). This occurred concomitant with an increase in neuroblasts (**Figure 7I**). Additionally, a one-hour pulse of EdU before euthanasia resulted in a higher proportion of EdU⁺ cells in cYAP^KO^/TAZ^HET^ *vs*. WT mice (**Figure 7J**). Overall, our data indicated that YAP/TAZ are required for quiescent NSC maintenance and its loss shifts the balance toward activation in the V-SVZ niche. Altogether, our data indicate that YAP/TAZ are crucial for generating the characteristic speckled structure of the V-SVZ ECM and for maintaining the essential interactions between qNSCs and these speckles, which are vital for preserving NSC quiescence.

## Discussion

Our study reveals a novel, niche-specific function of YAP/TAZ signaling in maintaining NSC quiescence in the adult V-SVZ by orchestrating the proper assembly of the ECM microenvironment. We show that NSCs in this niche initiate a quiescence program in response to BMP4, characterized by the production of a unique ECM proteome capable of independently inducing quiescence. This self-sustaining mechanism depends on YAP/TAZ-driven transcription, activated through NSC adhesion to the ECM generated in response to BMP signaling. In turn, YAP/TAZ promote further ECM remodeling, including deposition of fibrillar collagens, creating a feedback loop that maintains quiescence without requiring ongoing BMP4 exposure. Our findings suggest that a transient soluble signal can initiate the formation of a stable physical niche that preserves NSC quiescence.

Previous analyses have highlighted that the ECM of the V-SVZ is notably distinct from that of the cerebral cortex^24^ and that interactions between LMs and integrins in NSCs appear required for the proper assembly of speckles^30^. However, its functional contribution to niche dynamics and regulatory mechanisms remains poorly understood. Here, we address this gap by showing that BMP4 not only influences NSC state, but also actively induces the deposition of a matrix that closely reflects the *in vivo* ECM signature of the V-SVZ, including niche-specific matrisome components such as Anxa and S100 family proteins. This indicates that transient BMP4 signaling can induce ECM remodeling, establishing long-lasting structural conditions within the niche. A particularly compelling finding is that BMP4-induced ECM modifications are sufficient to induce quiescence in otherwise active NSCs *in vitro*, even in the absence of BMP signaling. This suggests a self-reinforcing mechanism in which quiescent NSCs actively generate a matrix that sustains their own quiescent state. YAP/TAZ signaling plays a central role in this process, linking ECM adhesion and mechanosensing to transcriptional programs that preserve niche organization. In this context, our iQ assay provides a versatile platform to explore adhesion-dependent mechanisms in NSC activation-to-quiescence transitions, enabling detailed studies of ECM-mediated regulation *in vitro*.

Further investigation may be needed to assess BMP-mediated ECM deposition *in vivo*, and how it is related to NSC dynamics in the V-SVZ niche. BMPs have garnered much interest since its establishment as a way of modeling a reversible quiescent-like state in hippocampal NSCs^11,64^. However, the precise mechanisms by which BMPs regulate this state remain poorly understood. In the SGZ, BMPs maintain NSC quiescence, preventing premature depletion^11,81,82^. Their role in the V-SVZ, however, is less clear. BMP diffusion is limited by ECM binding, particularly to heparan sulfates^32,34^ and the ependyma secretes BMP inhibitors like Noggin and LRP2, creating a gradient of BMP inhibition^83–85^. Increased BMP signaling in the V-SVZ reduces proliferation and neurogenesis, effects reversed by Noggin^83^ and reduced LRP2 increases Nestin^+^/Sox2^+^ NSCs with nuclear phospho-SMAD and Id3, while reducing Dlx2^+^ progenitors^84^. However, Smad4 deletion in GLAST^+^ cells skews differentiation toward oligodendrocytes^86^, highlighting potential cell type–specific effects. Our proteomic analysis of BMP4-induced qNSCs reveals that this morphogen plays a crucial role in shaping the V-SVZ ECM niche and modulates the acquisition of a specific quiescent matrisome signature that is shared by qNSCs *in vivo*.

Mechanotransduction via YAP/TAZ is classically associated with promoting proliferation, tissue regeneration, and stemness across diverse tissues^45,47,79^, including the developing and adult nervous system^49–53^. However, our findings uncover a context in which YAP/TAZ function conversely to preserve dormancy also through TEAD-mediated transcription. We show that either constitutive YAP activity or its activation in response to NSC-generated ECM *in vitro* is sufficient to induce a quiescent state. Besides, YAP/TAZ activity *in vivo* is required for maintaining normal ECM niche integrity. Loss of this organization leads to aberrant physical interactions, mislocalization of NSCs, and a shift towards activation, ultimately disrupting the balance between quiescent and active states. These results position YAP/TAZ as central mediators of the NSC-ECM feedback loop essential for sustaining a functional quiescent niche. This reveals an unprecedented role for these transcriptional coactivators always associated with proliferation and growth. Notably, the role of YAP/TAZ in adult NSC quiescence is unique to the V-SVZ, as evidenced by the role of YAP in NSC activation in the SGZ^53^. This discrepancy underscores the context-dependent nature of YAP/TAZ signaling, likely shaped by the distinct composition and structural organization of the ECM in each neurogenic niche. In the V-SVZ, the presence of specialized matrix structures, such as fractones^27,28,30,31,35^, and a unique matrisome profile^24^ may create a permissive environment for YAP/TAZ activation in quiescent NSCs. YAP/TAZ are well-known integrators of mechanical and ECM signals: they are activated by substrate stiffness and cytoskeletal tension and can themselves promote ECM remodeling through regulation of focal adhesion assemblies and matrix deposition^69,87,88^. For example, forced YAP expression can reprogram differentiated cells into tissue-specific stem cells^68^, and during colonic epithelium repair and YAP/TAZ are activated by ECM cues to induce a fetal-like regenerative state^89^. These findings highlight the remarkable plasticity of YAP/TAZ responses and support a model in which their function is tightly dictated by microenvironmental context. Beyond the V-SVZ, YAP/TAZ involvement in cellular dormancy has only been described in pathological settings, such as in lung cancer cells adapting to targeted therapy^90^.

Our results identify YAP/TAZ as essential regulators of fractone formation, providing the first molecular link between transcriptional regulation and these unique niche structures *in vivo*. These findings point to a broader conceptual shift, from quiescence as a state passively maintained by extrinsic cues to a dynamic, self-organized process in which qNSCs construct and maintain a specialized microenvironment that reinforces their own dormancy. Thus, transient signals like BMP4 may serve as initial instructive cues that, through YAP/TAZ-dependent pathways, lead to durable, ECM-mediated stabilization of the quiescent state.

## Acknowledgements

We acknowledge the support of the Servicio Central de Soporte a la Investigación Experimental (SCSIE-UVEG) and the Proteomics Facility at Universitat de València. We thank Maria Jose Palop for the technical support and Ana Domingo-Muelas for her support with the graphical design. We thank Drs. Enric Poch from Universidad CEU Cardenal Herrera (Valencia, Spain) and Jorge Oliver de la Cruz from Institute for Bioengineering of Catalonia (Barcelona, Spain) for the generous donations of plasmids for the study of ROCK and YAP function, respectively. During the preparation of this work we used ChatGPT in order to improve language and readability in some paragraphs. After using this tool/service, we reviewed and edited the content as needed and, therefore, we take full responsibility for the content of the publication. This work was supported by grants PID2020-117937GB-I00, RED2018-102723-T, CB06/05/0086 (CIBERNED) from Ministerio de Ciencia e Innovación (MICINN), Prometeo 2021/028 from Generalitat Valenciana, and ERC Advanced Grant 101098241 from the European Commission to I.F and by grants PID2020-114227RB-100 and CNS2022135758 from MICINN to C.G-S. P.D-A. and C.S. were recipients of FPI predoctoral contracts from the Ministerio de Ciencia e Innovación (MICINN), P.G-B is a recipient of a predoctoral contract from the Generalitat Valenciana and L.B-C. was a recipient of a FPU predoctoral contract from Ministerio de Educación, Cultura y Deporte (MECD).

## Author contributions

Conceptualization, L.B-C., I.F.; Methodology, L.B-C., P.G-B., P.D-A., JM.M-R., S.S., C.S., A.J-P., I.M-W., C.G-S.; Investigation, L.B-C., P.D-A., P.G-B., JM.M-R., S.S., C.S., A.J-P., I.M-W., C.G-S.; Writing – Original Draft, L.B-C., I.F.; Writing – Review & Editing, L.B-C., I.F., JM.M-R.; Visualization, L.B-C., P.D-A., JM.M-R.; Supervision, I.F.; Project Administration, I.F.; Funding Acquisition: I.F.

## Competing financial interest statement

The authors declare no competing financial interests.

## Methods

### Mouse strains and handling

Adult C57BL/6J wild-type (WT) mice (stock 000664), *Wwtr1^tm1.2Hmc^ Yap1^tm1.2Hmc^/WranJ* (strain 030532) and *6.Cg-Tg(Gfap-cre)73.12Mvs/J* (strain 012886) were obtained from the Jackson Laboratory and bred at the animal housing facility of the Universitat de València (Servei Central de Suport a la Investigació Experimental, SCSIE, Burjassot). YAP/TAZ mutant mice were obtained by crossing male double floxed *Yap/Taz* mice, which possess loxP sites flanking exon 2 of both the *Yap1* and *Wwtr1* (*Taz*) genes, with female GFAP-Cre mice. Housing and experimental procedures complied with the European Union 2010/63/UE directive and Spanish RD-53/2013 guidelines, under the supervision of an authorized veterinarian. Both males and females were used in all the experiments. Genetically modified mice were genotyped by end-point Polymerase Chain Reaction (PCR) of genomic DNA (gDNA) extracted from an ear punch using the Phire Animal Tissue Direct PCR kit (Thermo Fisher, cat. no. F140WH). For all *in vivo* and *in vitro* experiments, adult mice ranging from 2-4 months of age were used. For proliferation studies, one intraperitoneal (i.p.) injection of EdU (50_mg/kg of body weight) (Life Technologies, E10187) was injected 1_h prior to euthanasia (EdU-1h). For studying label retaining cells (LRCs), mice were i.p. injected with 50_mg/kg of BrdU (Sigma, B5002) every 1.5_h for 12_h (7 injections) and euthanized 21 days after the last injection.

### In utero electroporation (IUE)

IUE was used to introduce episomal plasmids encoding fluorescent proteins (mRFP or GFP) under the control of the constitutive CAG promoter into the lateral ventricles of C57BL/6 mouse embryos at embryonic (E) day E15.5 as previously reported^57^. Briefly, pregnant mice were deeply anesthetized with 2.5% (v/v) isoflurane at 0.8 ml/min and injected with 0.1 mg/kg of buprenorphine. Plasmid DNA (1-2 µg/μl) with 1/20 volume of 1% Fast Green (Sigma-Aldrich, F7252) was injected into the lateral ventricles of the embryos. Then, forceps-type platinum electrodes (Platinum Tweezertrode, 5 mm Diameter; BTX) were laterally placed around the head of the injected embryo and oriented with the positive electrode contacting the ventrolateral region of the injected hemisphere using a square wave electroporator (ECM 830 Square Wave Electroporation System, Btx). The electroporation conditions involved 5 pulses of 45 V of 80 ms, spaced 950 ms. Animals were maintained on the heating pad until recovery with additional analgesia and topically applied antibiotics to prevent infection. Brains from electroporated mice were analyzed by flow cytometry or processed for histological analysis at 2 months of age.

### Immunofluorescence

Young adult mice were anesthetized with an i.p. of medetomidine (0.5–1 mg/kg) and ketamine (50–75 mg/kg) in saline. Animals were perfused at 5.5 ml/min with 28 ml of saline followed by 83 ml of 4% paraformaldehyde (PFA) in 0.1M phosphate buffer (PB). Brains were post-fixed in PFA for 1 h at room temperature (RT), washed in PBS, and embedded in 4% agar for vibratome sectioning. Coronal sections (40-50 μm) were obtained using a Leica VT1000 vibratome and stored in sodium azide 0.05%. For whole-mount V-SVZ preparations, non-perfused animals were dissected fresh and fixed by immersion in 4% PFA with 0.5% Triton-X-100 for 1 h.

Samples were permeabilized and blocked with 10% horse serum and 0.1–0.2% Triton X-100 in PBS for 1 h at RT, followed by overnight incubation with the following primary antibodies at 4 °C: chicken anti-GFAP (1:800, Millipore, AB5541), chicken anti-GFP antibody (1:500, Aveslab, GFP-1020), rabbit anti-LM (1:500, Novus, NB300-144), mouse anti-β-catenin (1:400, BD, 610153), rabbit anti-β-catenin (1:100, Cell Signalling, 9587S), rat anti-Ki67 (1:1000, Invitrogen, 14569882), rabbit anti-aYAP (1:100, Abcam, ab205270), mouse anti-Arl13b (1:300, UCDavis, 75-287). After washing, samples were incubated with fluorescence-conjugated secondary antibodies for 1 h at RT in the dark: donkey anti-chicken AlexaFluor 488 (1:800, Jackson), donkey anti-chicken Cy3-conjugated AffiniPure (1:800, Jackson), donkey anti-rabbit AlexaFluor488 (1:800, Jackson), donkey anti-rabbit AlexaFluor Cy3 (1:800, Jackson), donkey anti-rat Alexa Fluor 647 (1:800, Jackson), donkey anti-mouse Alexa Fluor 647 (1:800, Jackson). Nuclei were counterstained with 4′,6-diamidino-2-phenylindole (DAPI, 1 μg/ml) for 5 min and mounted using FluorSave™ for visualization and preservation. For BrdU detection, sections were pre-incubated in 2_N HCl for 20_min at 37_°C and neutralized in 0.1_M sodium borate (pH 8.5) prior to incubation with blocking solution. EdU detection was carried out using the Click-iT^TM^ EdU Alexa Fluor^TM^ 555 Imaging Kit (ThermoFisher, C10338). Images were acquired with an Olympus FV10i confocal microscope (405, 458, 488, and 633 nm lasers). For analyzing apico-basal positions *in vivo*, images were acquired using a super-resolution confocal microscope (Zeiss, LSM 980).

For *in vitro* cultures, cells were fixed (2% PFA, 15 min) and thoroughly washed prior to immunostainings. Cells were blocked with 10% horse serum and 0.1% Triton X-100 in PBS for 1 h at RT, followed by overnight incubation with rat anti-Ki67 (1:1000, Invitrogen, 14569882) and rabbit anti-aYAP (1:100, Abcam, ab205270) overnight at 4 °C:. After washing, cells were incubated with fluorescence-conjugated donkey anti-rabbit AlexaFluor488 (1:800, Jackson) and donkey anti-rat Alexa Fluor 647 (1:800, Jackson), combined with rhodamine phalloidin (1:400, Thermo Fisher, R415) for labelling F-actin. Images were acquired with an Olympus FV10i confocal microscope (405, 458, 488, and 633 nm lasers).

### Expansion microscopy and optical clearing

Expansion microscopy was performed as previously described^91^. Immunostained V-SVZ whole-mounts were incubated overnight at RT with Acryloyl-X SE (Thermo Fisher) to promote protein cross-linking, directly in MatTek dishes (P35G-1.5-14-C). Gelling solution was added for 30 min at 4_°C in the dark to ensure infiltration. After removing the solution, fresh gelling solution was added, and samples were overlaid with a 15_µm coverslip to form a flat gel. Polymerization was carried out for 2_h at 37_°C in the dark. Gels were then digested overnight with proteinase K. Expansion was achieved by replacing the digestion solution with distilled water (3–4 washes, 10–15_min each). Expanded gels were stabilized in 2% agarose, submerged in water, and imaged using an inverted confocal microscope with a water-immersion objective (Leica, TCS SP8). For enhanced imaging resolution and fine cell interaction analysis, V-SVZ wholemounts were optically cleared using the X-Clarity™ system (Logos Biosystems, C3000). For optically cleared tissue, immunostainings were performed using the DeepLabel Antibody Kit (Logos Biosystem, C33001), following the manufacturer’s instructions. Then, sections were mounted in X-CLARITY mounting solution for refractive index homogenization and analyzed using a super-resolution confocal microscope (ZEISS, LCS 980).

### Transmission electron microscopy

Electroporated adult mice were anesthetized and perfused with 4% PFA and 0.5% glutaraldehyde (EM grade, Electron Microscopy Science) in PB before obtaining 50 μm vibratome sections. For the identification of IUE-stained cells, sections were blocked and incubated with a primary chicken anti-GFP antibody (1:200, Aves Labs) for 48 h at 4 °C with agitation, followed by a secondary antibody conjugated to ultrasmall colloidal gold (1:50, Aurion). Silver enhancement (Aurion R-Gent SE-LM) was performed for 20 min in the dark to enlarge gold particles, followed by gold toning using 0.05% gold chloride and 0.3% sodium thiosulfate. Reacted sections were osmicated (1% OsO4 in PB, 20 min), dehydrated in graded alcohols to propylene oxide, and plastic-embedded flat in Durcupan (Sigma). To study selected IUE-stained qNSCs, serial 1.5 µm-sections were cut with a diamond knife and stained with 1% toluidine blue. Subsequently, the area of interest was trimmed, and ultrathin sections (50-70 nm) were obtained, stained with lead citrate, and examined in a Jeol JEM 1010 electron microscope. Adult cYAP^KO^/TAZ^HET^ mutants and WT mice were transcardially perfused with saline followed by 2% PFA and 2.5% glutaraldehyde in PB. 200 μm coronal sections were post-fixed in 2% osmium tetroxide for 2 h, rinsed, dehydrated in graded alcohols to propylene oxide and embedded in Durcupan. Ultrathin sections containing the V-SVZ were obtained, stained with lead citrate, and examined in a Jeol JEM 1010 electron microscope.

### Image analysis

To quantify speckle number, size and interaction with GFAP^+^ cells, confocal images of whole-mount V-SVZ preparations stained for GFAP and LM were analyzed using a custom Fiji macro^92^. Max intensity z-projections of planes containing speckles were generated. GFAP channel was segmented by Li thresholding after contrast enhancement (saturated = 0.5). Objects smaller than 2 pixels were excluded to obtain the refined binary masks. LM channel was preprocessed by contrast enhancement (saturated = 0.4, normalize) and background subtraction (rolling = 100) before binarization based on thresholding (>42; 8-bit). The resulting mask was refined by removing objects not fitting size (1.5-120 µm^2^) and circularity (0.1-1.00) criteria, filtering out vasculature. This speckles mask was used to measure speckle number and size, and applied to the GFAP mask to calculate the GFAP-speckle contact area, which was then normalized to the total GFAP^+^ area to determine the interaction percentage. For each animal (n = 4), a minimum of 8 z-stacks were analyzed.

GFAP⁺ BrdU⁺ label-retaining cells (LRCs) were quantified in V-SVZ whole-mounts. For each animal, a minimum of 9 fields were analyzed, covering anatomically comparable rostral, medial, and caudal regions in both WT and cYAP^KO^/TAZ^HET^ samples. Quantifications were performed on z-stack projections (16_µm total thickness) by counting the number of GFAP⁺ BrdU⁺ cells per field (7,2 x 10^-4^ mm^3^) using the *FluoView* software (Olympus). To assess the distribution of GFAP labeling along the apico-basal axis, whole-mount V-SVZ preparations immunostained for GFAP and β-catenin, quantification was carried out using a custom Fiji macro. Ependymal surface was defined as the reference plane (0 µm) and GFAP^+^ regions were segmented across successive optical sections by thresholding (>195; 12-bit) and refined excluding objects < 3 µm^2^. GFAP^+^ area was measured along the different optical sections and then normalized to the total GFAP^+^ area for each animal, yielding a standardized distribution profile. A minimum of 6 z-stacks were analyzed per animal. Depth color-coded images were generated from whole-mount preparations immunostained for β-catenin and GFAP. Image stacks consisting of 60 optical sections at 0.17_µm intervals (total depth = 10.2_µm) were processed for visualization. Z-stacks were imported into Fiji (ImageJ, NIH) and converted to temporal color-coded hyperstacks using the Temporal-Color Code function. Depth-based color coding was applied using the “16 colors” lookup table, allowing clear visual distinction of structures along the z-axis. The ependymal layer was identified based on β-catenin staining and used as the positional reference (depth = 0_µm).

To quantify nuclear YAP intensity *in vivo*, z-stacks (3_µm total thickness from the ependymal surface) were acquired from anatomically matched regions of V-SVZ whole-mounts from WT and cYAP^KO^/TAZ^HET^ mice (n = 4 mice *per* group). GFAP⁺Ki67⁺ and GFAP⁺Ki67⁻ nuclei boundaries were manually delineated based on DAPI staining using the freehand selection tool in Fiji (ImageJ, NIH), and mean nuclear YAP fluorescence intensity was measured for each cell nucleus. All imaging settings were kept constant throughout image acquisition. For each experimental group, the average nuclear YAP intensity was calculated across all quantified cells for statistical analysis. For YAP nuclear intensity quantification *in vitro*, 12-bit confocal z-stacks of cells stained for DAPI, Ki67, and YAP were processed using a custom Fiji macro. Sum intensity z-projections were generated for each channel. Nuclei were segmented from the DAPI projection: images were preprocessed by background subtraction (rolling = 25) and median filtering (radius = 2) and thresholding was performed using the RenyiEntropy method, followed by binarization and watershed-based separation of overlapping nuclei. Finally, objects smaller than 25 pixels were excluded. The resulting nuclear ROIs were applied to the Ki67 and YAP projections to measure mean nuclear intensity. A minimum of 50 cells per culture were scored establishing a >250 intensity threshold to be classified as Ki67 positive.

### Neurogenic lineage phenotyping and cell sorting

We used a flow cytometry-based approach to characterize V-SVZ neurogenic lineage cells, as described previously^6,59^. V-SVZ tissue from 2–4-month-old mice was enzymatically and mechanically dissociated into single-cell suspensions using the Neural Tissue Dissociation Kit (T) with an automatic GentleMACS Dissociator. Cells were filtered, centrifuged (300*g*, 10 min), and incubated with lineage-specific antibodies in flow cytometry buffer (0.1% Glucose, 10_mM HEPES, 2_mM EDTA and 0.5% BSA in HBSS) for 30 min at 4 °C: CD45-BV421 or CD45-BUV395 (1:100, BD), O4-AF405 (1:50, R&D), CD31-BUV395 (1:100, BD) or CD31-BV421 (1:100, BD), Ter119-BUV395 or Ter119-BV421 (1:200, BD), EGF-Alexa488 or EGF-Alexa647 (1:300, Molecular Probes), CD24-PerCP-Cy5.5 (1:300, BD) or CD24-BB700 (with EdU detection; 1:300; BD), GLAST-PE (1:50, Miltenyi) or GLAST-APC (with EdU detection; 1:50, Miltenyi), and CD9-APC-Vio770 (1:20, Miltenyi). Then, samples were washed in the same buffer, filtered, counterstained with DAPI (0.1 μg/ml) to exclude dead cells and analyzed using a LSR-Fortessa cytometer (Becton Dickinson) with 350, 405, 488, 561 and 640 nm lasers. Briefly, microglia, oligodendrocytes, erythrocytes, and endothelial cells, named as Lin^+^, were first excluded using specific markers conjugated to the same fluorochrome. Then, Lin^-^ cells were stratified into: (1) neuroblasts (NBs: GLAST^-^/CD24^high^) that are further separated into: NB1 (EGFR^+^) and NB2 (EGFR^-/low^); (2) neural progenitor cells (NPCs) comprised of two different subpopulations: NPC1 (GLAST^-^/CD24^-/low^) and NPC2 (GLAST^+^/CD24^high^), both EGFR+; (3) non-neurogenic astrocytes (GLAST^+^/CD24^-/low^/CD9^low^), and (4) neural stem cells (NSCs: GLAST^+^/CD24^-/low^/CD9^high^) which are further divided into quiescent (qNSCs: GLAST^high^/ EGFR^-/low^), primed-for-activation (pNSCs: GLAST^low^/EGFR^-/low^) and activated (aNSCs: GLAST^low^/EGFR^+^)^59^. To evaluate proliferation in EdU-injected mice, samples were fixed with 100_µl of Cytofix/Cytoperm™ solution (BD Biosciences, cat. no. 554722) diluted 1:4, for 20 minutes in the dark at 4_°C and processed using the Click-iT™ Plus EdU Alexa Fluor™ 555 Flow Cytometry Assay Kit (ThermoFisher, C10638) according to manufacturer’s instructions. For LM detection by flow cytometry, cells were mechanically disaggregated on ice and LM primary (1:100, Novus, NB300-144) and secondary anti-rabbit Alexa Fluor® 647 (1;400, Molecular Probes, A31573) antibodies were combined with the marker panel for the neurogenic cell populations. LM was assessed by evaluating the median fluorescence intensity (MFI) from each population and removing unspecific background signals by using fluorescence minus one (FMO) controls.

For aNSC cell sorting, the V-SVZ of two C57BL/6J mice were dissected, pooled, and dissociated as previously described. Cells were stained with our flow cytometry panel and BD FACSAriaTM Fusion Flow Cytometer with 350, 405, 488, 561 and 640 nm lasers was used for cell sorting. p48-well culture dishes (1 cm²) were coated with cMtx or bMtx, supplemented with the BMP4 antagonist Noggin (RyD, 1967) at 0.4 μg/ml to neutralize residual BMP4. Coated wells were rinsed with distilled water and filled with 500 μl of growth medium supplemented with bFGF and EGF. A total of 300 aNSCs were directly sorted into each well and incubated at 37_°C with 5% CO₂. Cells were manually counted at 24, 48, and 72 h to evaluate division. Viable cells were identified as refringent and quantified as singlets or aggregates (doublets or triplets), expressed as a percentage of total live sorted cells.

### Neurosphere culture and subculture

Primary neurosphere cultures were performed as previously reported^93^. V-SVZ were dissected and minced in sterile PBS and digested for 30 min at 37 °C with a pre-activated papain solution (12 U/ml papain in EBSS with 0.2 mg/mL L-cysteine and 0.2 mg/ml EDTA), followed by mechanical dissociation. The resulting cell suspension was washed, centrifuged, and resuspended in NSC growth medium (5mM HEPES, 0.1% sodium bicarbonate, 2mM glutamine, 1X antibiotic/antimycotic, 0.7U/ml heparin and Hormone Mix in DMEM/F12) containing 20 ng/ml EGF (Gibco, 53003-018) and 10 ng/ml bFGF (Sigma, F0291). The suspension was seeded in p48-well plates (500 μl/well) and cultured for 7–10 days to enrich NSCs and progenitors, which grew as floating neurospheres.

For NSC maintenance, cells were subcultured every 5 days, and neurospheres were collected, centrifuged (130*g*, 7 min), and enzymatically digested with Accutase® solution (Sigma, A6964) for 10 min at RT followed by mechanical disaggregation. Then, cells were washed, centrifuged, and resuspended in growth media. Cells were seeded at a density of 10,000 cells/cm² for continued maintenance. For transfection experiments, supplement 1X B27 was added (Thermo Fisher, 12587010). For neurosphere formation assays, cells were seeded at *pseudoclonal* density (5 cells/μl) and kept in the incubator at 37°C 5% CO2 for 7 DIV to allow neurosphere formation. When required, cells were seeded at the same density under adherent conditions using commercial Matrigel (200_μl/cm²; Corning™, 356234), which was incubated overnight and thoroughly washed with distilled water prior to cell seeding. Phase-contrast images were acquired using a Nikon ECLIPSE TE2000-S inverted microscope with phase contrast optics. Numbers of neurospheres were counted manually and cells photographed for sphere diameter manual evaluation using Fiji software.

### NSC *in vitro* treatments and proliferation assessment

To assess proliferation in NSC cultures, cell tracer DFFDA was used as reported^6^. Single-cell suspensions (1 million cells) were washed with PBS, centrifuged, and incubated with 2 μg/ml DFFDA (CellTraceTM Oregon Green 488 Carboxy-DFFDA-SE, C34555) in PBS for 7 minutes at 37 °C in a shaking thermoblock and protected from light. Then, cells were seeded in 20 ng/ml EGF and 10 ng/ml bFGF supplemented growth medium in uncoated or coated wells and DFFDA dilution was assessed by flow cytometry after 4 DIV. To do so, we defined a slow-cycling population (DFFDA^high^) as the top 10% of cells with highest DFFDA intensity^6^. Experimental conditions were compared to the reference DFFDA^high^ population from the control group. For quiescence induction *in vitro*, cells were cultured in uncoated wells and treated with 50 ng/ml BMP4 (RYD, 314-BP) or its vehicle (4 mM HCl with 0.1 % bovine serum albumin (BSA)) for 4 DIV in 20 ng/ml EGF and 10 ng/ml bFGF or in each mitogen alone. For ROCK inhibition, cells were cultured in growth medium with 25 μM Y27632 (Sigma, Y0503) or its vehicle (DMSO) for 4 DIV. Then, cells were detached by adding Accutase® to cell culture plates, centrifuged and incubated with mouse EGF–A647 (1:300, Molecular Probes, E35351) 20 min on ice and protected from light to assess EGFR levels by flow cytometry. Alternatively, cells were incubated with 10 μM EdU (ThermoFisher, C10338) for 1 h at 37 °C in growth medium. Following incubation, cells were collected, dissociated using Accutase®, and fixed and permeabilized with Cytofix/Cytoperm (ThermoFisher, BD 555028) 20 min at 4 °C and protected from light. Then, Click-iT® EdU Alexa Fluor™ 555/647 Imaging Kit (ThermoFisher, C10338) was used for EdU detection.

### iQ assay

For the induced-quiescence (iQ) assay, neurosphere cell suspensions were seeded in growth medium with 50 ng/ml BMP4 or vehicle (4 mM HCl, 0.1% BSA) for 4 DIV. Then, conditioned media (CM) was collected and centrifuged to remove debris. When not used immediately, CM was stored at –80_°C. To neutralize residual BMP4 activity, 0.4 μg/ml Noggin (RyD, 1967) was added to the CM prior to its use. Control and BMP4-CM were used to coat cell culture well plates (200 μl/cm²) overnight at 37 °C. Before seeding, derived BMP4-CM matrix (bMtx) or control-CM matrix (cMtx) were removed, and wells washed several times with distilled water. In specific experiments, plates were pre-coated with Matrigel (CorningTM, 356234) (200 μl/cm²) overnight, followed by washes and subsequent coating with bMtx or cMtx. Dissociated neurosphere cells were seeded onto the prepared matrices in growth medium supplemented with bFGF and EGF. For adhesion analysis, cells were seeded at 10,000 cells/well in 48-well plates and imaged using an Incucyte® S3 microscope under stable conditions of temperature, humidity, and CO_2_ concentration. Label-free, phase contrast images were acquired using a 20X objective. Cell segmentation was performed using a custom trained CellPose model accessible through Zenodo (doi: 10.5281/zenodo.15623212). Ground truth masks were generated with manual annotation in QuPath^94^, and segmentation and shape quantification were carried out using the QuPath–CellPose extension^95^. Since floating cells display a rounded morphology but may acquire a range of morphological phenotypes once adhered, cells with a circularity < 0.5 were classified as adhered. For evaluating proliferation, cells were loaded with cell tracer DFFDA (CellTraceTM Oregon Green 488 Carboxy-DFFDA-SE, C34555) prior to the seeding and, after 4 DIV, cells were collected, incubated with EGF-647 (1:300, Molecular Probes, E35351) 20 min on ice and protected from light, and analyzed by flow cytometry. For detachment and disaggregation, Accutase® (Sigma, A6964) was applied directly to adherent cultures or to pelleted neurospheres.

### NSC transfections

NSCs were transfected with plasmids to disturb specific proteins. The ROCK1 WT and dominant negative ROCK1 Δ5 constructs were kindly supplied by Enric Poch from Universidad CEU Cardenal Herrera (Valencia, Spain). For evaluating YAP function and YAP-TEAD activity, the following plasmids were kindly provided by Jorge Oliver de la Cruz from Institute for Bioengineering of Catalonia (Barcelona): pBabe-YAP-5SA (constitutively active YAP), pBabe-YAP-5SA/S94A (constitutively active, TEAD-binding deficient YAP construct), pBabe (control, empty vector) and reporter pLL3.7 FLAG-YAP1-TEAD-P-H2B-mCherry. ShYAP plasmid was obtained from Addgene (42540). Plasmids were amplified, purified, and confirmed by restriction enzyme digestion. Primary NSCs (4-5 million cells *per* nucleofection) were resuspended in 90 μl of the nucleofection buffer containing 4 μg of the plasmid of interest and using Lonza nucleofector kit (vpg-1004) and Nucleofector 2b device with G-013 preset program. Cells were seeded in NSC groth medium supplemented with bFGF and EGF and with 1X B27 for recovery (Thermofisher, 12587010). YAP reporter activity was evaluated as mCherry mean fluorescence intensity (MFI) normalized to GFP MFI among GFP^+^ population, by flow cytometry 48 h post-seeding. For YAP gain-of-function experiments and silencing with shRNA, cells were co-transfected with pEGFP.N1 plasmid (4:1). After a 24 h recovery, cells were disaggregated, washed with Debris Removal Solution (Miltenyi, 130-109-398), counted and seeded at normal density (10,000 cells/cm²) in NSC medium with B27 supplement and with EGF and bFGF. Experiments were performed 24 h after reseeding, except when treating with BMP4, for which cells were cultured for 48 h. Cell proliferation was assessed among the transfected (GFP^+^) population.

### Quantitative PCR

RNA was extracted using the RNeasy Micro or Mini kit (Qiagen, 74.134) and quantified with the Qubit™ RNA HS Assay Kit (Thermo Fisher, 32852). For reverse transcription, 0.5–1 μg of RNA was converted to cDNA using the PrimeScript RT kit (Takara, RR037A) with random hexamers and oligo-dT primers. Gene expression was analyzed by qPCR using TaqMan™ probes and Premix Ex Taq™ Kit (Takara, RR390) in a Step One Plus system (Applied Biosystems). A standard 40 cycle program with annealing and extension steps at 60 °C was used. Relative expression was calculated with the 2^- ΔΔCt^ method, normalized to Gapdh and 18S geometric mean.

### LM immunoprecipitation and Western blot

To assess the presence of LM in NSC control and BMP4-CM, immunoprecipitation (IP) was performed, followed by Western blotting (WB), as high BSA levels in the CM interfered with direct protein detection. For IP, 1 ml of CM of each condition (n = 4 independent cultures) was incubated overnight at 4_°C on an orbital shaker with 5_μg of anti-LM antibody (Novus Biotechne, NB300-144) or an isotype control. Samples were then incubated with 50_μl of magnetic Protein G Dynabeads (Invitrogen, 10003D) for 1_h at RT on a rotating wheel. After four PBS washes using a magnetic separator (DynaMag, 12321D), IP complexes were eluted in 50_μl of Laemmli buffer by heating at 95_°C for 5_min. Samples were separated on 7% SDS-PAGE gels and transferred onto nitrocellulose membranes using the Mini Trans-Blot® Cell system (Bio-Rad). Membranes were blocked with 5% skimmed milk in TBS-T for 1_h at RT and incubated overnight at 4_°C with anti-LM antibody (2_μg/ml; 1:500, Novus, NB300-144) diluted in TBS-T with 5% BSA. Despite using the same antibody for IP and detection, LM was distinguishable from antibody light and heavy chains based on molecular weight. After TBS-T washes, membranes were incubated with HRP-conjugated anti rabbit secondary antibodies (1:5000, Dako, P0449) for 1_h at RT. Signal was developed using SuperSignal™ West Femto substrate (Thermo Scientific, 34095) and imaged with a Mini HD 9 chemiluminescence system (Uvitec).

### Quantitative proteomic profiling

NSC cultures (n = 4) were treated with BMP4 (50_ng/ml) or vehicle (4 mM HCl with 0.1 % BSA) in NSC growth medium supplemented with EGF and bFGF for 4 DIV, cells were collected by centrifugation (400*g*, 10_min) and pellets were processed by the Proteomics Facility at Universitat de València. Proteins were extracted and digested with Lys-C/trypsin using the EasyPep MS Sample Prep Kit (Thermo Scientific). Peptides were desalted, dried, and resuspended in 2% acetonitrile/0.1% TFA. LC–MS/MS analysis was performed on an ekspert nanoLC 425 system coupled to 6600plus TripleTOF mass spectrometer (SCIEX), using DIA (60-min SWATH acquisition, 100 variable windows). SWATH data were analyzed with DIA-NN 1.8 (Data-Independent Acquisition by Neural Networks). Automatic inference was set for mass accuracy. Any LC (High accuracy) was used as quantification strategy. Protein was inferred at gene level and output was filtered at 0.01 FDR. The mass spectrometry proteomics data have been deposited to the ProteomeXchange Consortium via the PRIDE^96^ partner repository with the dataset identifier PXD064825.

NSC cultures (n = 3) were treated with BMP4/vehicle for 4 DIV in growth medium without BSA. Then, CM was collected, mixed to reduce variability, and used to coat cell culture wells overnight. The following day, conditioned media was removed and the coatings (cMtx and bMtx) were washed and lysed in 1.5_ml of 50_mM ammonium bicarbonate containing 1_µg of Lys-C/trypsin (ThermoFisher) and digested overnight at 37_°C with gentle shaking, without prior reduction or alkylation. Digested peptides were acidified to 1% TFA, concentrated to 50_µl, and quantified with Qubit (Invitrogen). 200_ng of peptides were diluted in 0.1% formic acid, loaded into Evotip Pure tips (Evosep), and analyzed on a timsTOF fleX mass spectrometer (Bruker) using the Evosep One system (30 SPD method) in ddaPASEF mode. Protein identification was performed using MSFragger via FragPipe using default settings against the SwissProt database (FDR ≤1%). Data are available via ProteomeXchange with identifier PXD064830.

NSC cultures (n = 4) were transfected with YAP-5SA, YAP-5SA/S94A or an empty (pBabe) vector together with an pEGFP.N1 in a 1:4 proportion. After 48 h, cells were sorted using a BD FACSAriaTM Fusion Flow Cytometer. Immediately after cell sorting, cells were pelleted, supernatants removed and 20_µl of 0.001% DDM in 50_mM ammonium bicarbonate (ABC) were added to each cell pellet. Samples were then processed again at the Proteomics Facility (Universitat de València). Lysates were agitated, sonicated, and heated at 95_°C for 20_min. Protein content was quantified (Qubit, Invitrogen), reduced (2_mM DTT, 1_h, 37_°C), alkylated (5.5_mM IAM, 30_min, RT), and digested overnight with Lys-C and trypsin (200_ng, Thermo Fisher). Peptides were acidified (1% TFA) and 200_ng were loaded into Evotip tips and separated on an Evosep One system (30 SPD gradient). MS analysis was performed on a TimsTOF fleX (Bruker) in diaPASEF mode. Data were analyzed with DIA-NN v1.8 (FragPipe v21.1) using an in silico–predicted *Mus musculus* spectral library. Proteins were quantified at 1% FDR and gene-level data were used for downstream analysis. Data are available via ProteomeXchange with identifier PXD064829.

### Proteomic analysis

Quantitative proteomics data were imported from a matrix generated using DIA-NN. Protein intensities were rounded to the nearest integer to be able to perform differential protein expression analysis using the DESeq2 R package (v1.44.0). The analysis was performed with a paired design formula that included both the treatment group (Group) and the pairing variable (Subject) to account for inter-individual variation (∼ Subject + Group). This allowed for within-subject comparisons between treatment conditions. Proteins were retained for analysis if they had an intensity value of at least 1 in two or more samples. The control group was set as the reference level for comparison. The DESeq2 pipeline was then executed with default normalization parameters to estimate size factors, dispersions, and model fitting. Principal component analysis (PCA) was used to evaluate the similarity of the biological replicates, and an MA plot was generated to visualize the relationship between mean protein expression and change in expression between the two conditions for all proteins after filtering and size factor calculations performed by DESeq2. False discovery rate-adjusted p-values < 0.05 were extracted to identify proteins differentially expressed between subject and control samples. The lists of up- and down-regulated proteins were used as inputs for functional enrichment analysis. The analyses were based on the use of the “enrichGO” function as implemented in the clusterProfiler R package (4.12.0). Functional enrichment analysis for Biological Process (BP) ontology categories was performed using mouse-specific GO annotations provided by the *org.Mm.eg.db* database. Enriched GO term redundancy was minimized by using the “simplify” function of the same package, applying the following parameters: cutoff = 0.5, by = “p.adjust”, select_fun = min, measure = “Wang”, semData = NULL. The GOcircle function from the GOplot R package (1.0.2) was used to generate circular plots of representative GO terms. Spectral counts were employed to determine protein presence or absence across the analyzed samples. To improve data quality and specificity, common contaminants were excluded, and protein entries were filtered to retain only those annotated as originating from Mus musculus or Rattus norvegicus, based on UniProt identifiers ending in ‘_MOUSE’ or ‘_RAT’. Proteins were further annotated by cross-referencing with the Matrisome Database to identify extracellular matrix components. The STRING database^97^ was used to generate a protein-protein interaction (PPI) network, and to evaluate the statistical significance of the number of network connections among the identified proteins in bMtx and cMtx. STRING’s built-in GO term analysis tool was used to analyse the functional enrichment of the members of the network. All heatmaps were generated using the ComplexHeatmap (v2.20.0) and ggplot2 (v3.5.1) packages in R. For expression heatmaps, the Spearman rank correlation distance was used as the base for the hierarchical grouping and clustering of genes, unless otherwise stated. The overlap between lists of genes was represented as Venn Diagrams, generated by the ggVenn package (0.1.10).

### Statistical analysis

All statistical tests were performed using the GraphPad Prism Software, version 9.0.1 for Mac (http://www.graphpad.com). Significant differences between paired groups were assessed using a paired two-tailed Student’s t-test. Repeated measures one-way ANOVA with Tukey post-hoc multiple comparisons test was performed when comparing three or more experimental conditions. Statistical analysis of GFAP area distribution among z-planes was performed using two-way ANOVA. When comparisons were carried out with relative values (ratios and percentages), data were first normalized with *arcsin* transformations. Statistical comparisons between groups in proteomics-related boxplot representations were performed using the ggpubr package in R (0.6.0), applying Wilcoxon rank-sum tests. p-values lower than 0.05 were considered as statistically significant and referred as *p<0.05, **p<0.01 and ***p<0.001. Data are always presented as the mean ± standard error of the mean (SEM). The number of experiments carried out with independent cultures/animals (n) is shown as dots in the graphs and listed in the Figure Legends.

## Notes

### Competing Interest Statement

The authors have declared no competing interest.

