## Supplementary material for "YAP/TAZ create a physical niche for the maintenance of adult neural stem cell quiescence": Video legends

### Supplementary Video Legends

**Supplementary Videos 1. Time-lapse imaging of NSCs cultured for 4 DIV on uncoated culture wells. A.** NSCs in growth medium supplemented with EGF and bFGF. **B.** NSCs in growth medium supplemented growth medium and BMP4 (50 ng/mL); cell adhesion becomes evident after 2 DIV.

**Supplementary Videos 2. Time-lapse imaging of NSCs cultured over 4 DIV in growth medium supplemented with EGF and FGF under different matrix conditions. A.** NSCs on control-CM matrix (cMtx), pre-coated onto wells overnight and thoroughly washed before seeding. **B.** NSCs on cMtx pre-incubated with the BMP4 antagonist Noggin (0.4 μg/ml) prior to coating. **C.** NSCs on BMP4-CM matrix (bMtx), also used to coat wells overnight and washed before cell seeding. **D.** NSCs on bMtx pre-treated with Noggin (0.4 μg/ml).

**Supplementary Videos 3. Time-lapse imaging of WT and cYAP$^{KO}$/TAZ$^{HET}$ NSC cultures over 4 DIV in growth medium supplemented with EGF and FGF under different matrix conditions. A.** WT cells cultured on control-CM matrix (cMtx), which was pre-coated onto wells overnight and thoroughly washed prior to seeding. **B.** cYAP$^{KO}$/TAZ$^{HET}$ cells cultured on cMtx **C.** WT cells cultured on BMP4-CM matrix (bMtx), similarly pre-coated and washed before seeding. **D.** cYAP$^{KO}$/TAZ$^{HET}$ NSCs cultured on bMtx.
