## Supplementary figures for "YAP/TAZ create a physical niche for the maintenance of adult neural stem cell quiescence"

[1]Instituto de Biotecnología y Biomedicina (BioTecMed), Universidad de Valencia, Spain.
[2]Departamento de Biología Celular, Biología Funcional y Antropología Física. Universidad de Valencia, Spain.
[3]Centro de Investigación Biomédica en Red sobre Enfermedades Neurodegenerativas (CIBERNED), Spain.
[4]Instituto de Biomedicina de Valencia, Consejo Superior de Investigaciones Científicas (IBV-CSIC), Valencia, Spain.

Running title: *YAP maintains stem cell quiescence*

Keywords: *ventricular-subventricular zone, NSC, extracellular matrix, BMP, YAP/TAZ, ROCK, matrisome*

---

### Supplementary figure 1

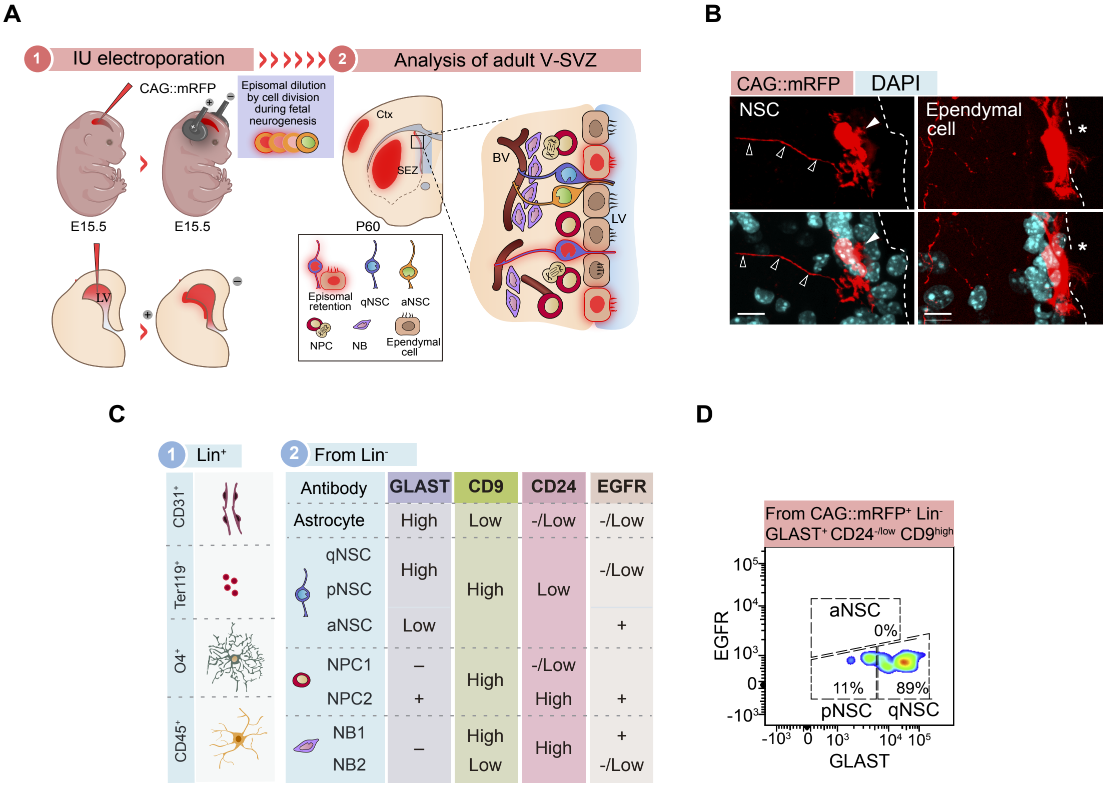


**Supplementary Figure 1. In utero (IU) electroporation to identify quiescent NSCs. A.** Schematic representation of the in utero electroporation strategy used to introduce CAG::mRFP plasmids into embryonic progenitors of the V-SVZ and assess episomal fluorescence retention in 2-month old mice. **B.** Representative confocal image showing mRFP expression in NSCs and ependymal cells following in utero electroporation. The asterisk marks multiciliated ependymal cells; the filled white arrowhead indicates a NSC primary cilium; empty arrowheads denote basal projections of NSCs. Dotted lines delimitate the lateral ventricle. **C.** Summary table of the multiparametric flow cytometry strategy used for neurogenic lineage characterization (adapted from Belenguer et al., 2021a). Relative expression levels of key surface markers used to identify distinct V-SVZ populations are indicated. **D.** Flow cytometry analysis of $RFP^{+}$ cells retaining episomal CAG::mRFP plasmids in the adult V-SVZ (2 months post-electroporation). EGFR levels (measured via fluorescently labeled EGF) and GLAST expression were used to define quiescent, primed, and activated NSC subpopulations. The gating strategy is outlined in the header. Scale bars:10 µm.

### Supplementary Figure 2

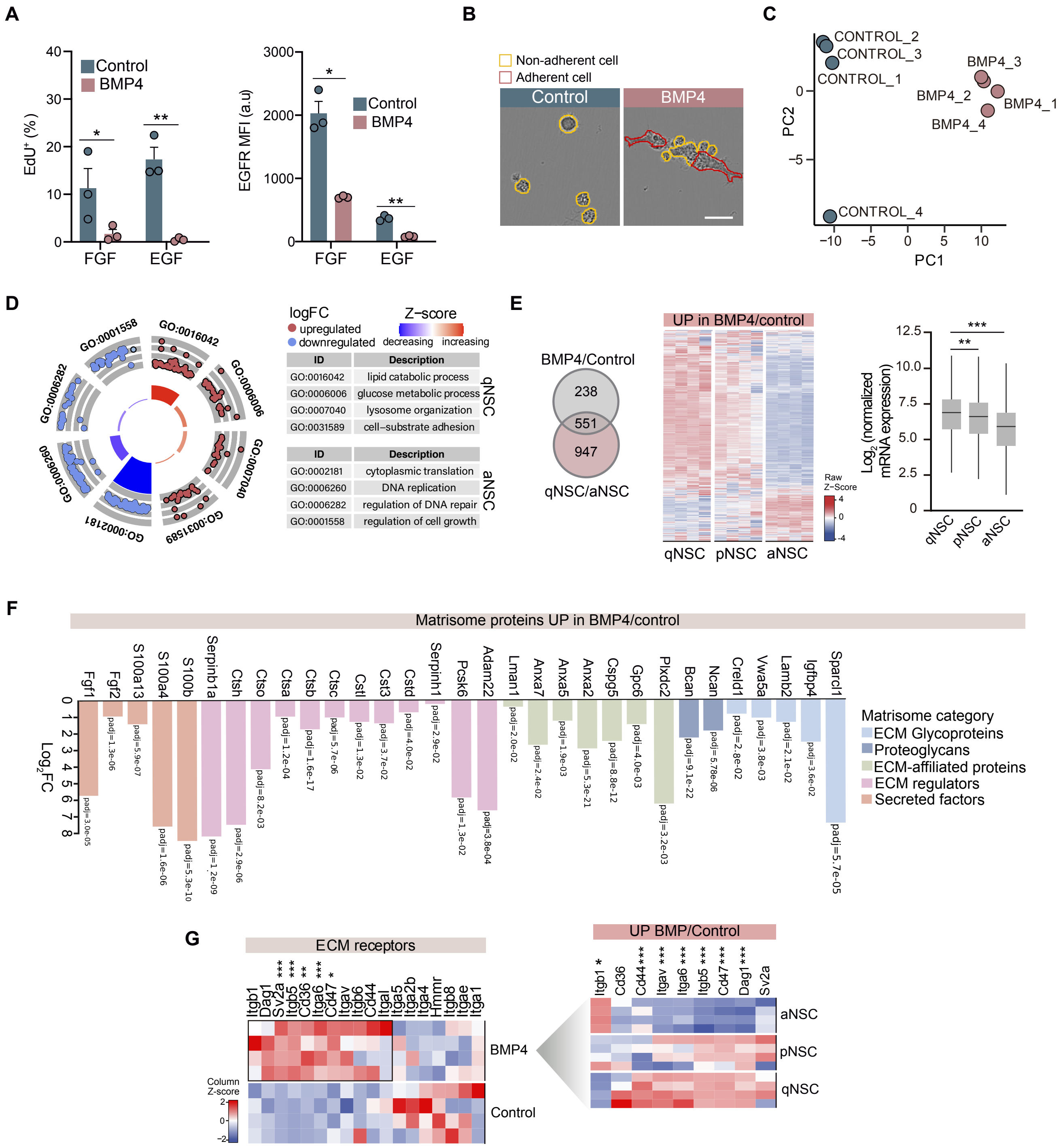


| ID | Description | |
|---|---|---|
| GO:0016042 | lipid catabolic process | qNSC |
| GO:0006006 | glucose metabolic process | qNSC |
| GO:0007040 | lysosome organization | qNSC |
| GO:0031589 | cell–substrate adhesion | qNSC |
| GO:0002181 | cytoplasmic translation | aNSC |
| GO:0006260 | DNA replication | aNSC |
| GO:0006282 | regulation of DNA repair | aNSC |
| GO:0001558 | regulation of cell growth | aNSC |

**Supplementary Figure 2. BMP4 signaling induces a quiescence-associated proteomic signature in NSCs. A.** Percentage of EdU⁺ cells (1-h pulse) and EGFR median fluorescence intensity (MFI) after 4 days in vitro (DIV) under mitogenic stimulation with either FGF or EGF, with or without the addition of BMP4 (50 ng/mL), assessed by flow cytometry. **B.** Phase contrast images with segmented adherent and non-adherent cells in control and BMP4 condition. **C.** Principal component analysis (PCA) of SWATH-MS proteomic profiles from untreated (control) and BMP4-treated NSCs. **D.** Circular plot summarizing four representative Gene Ontology (GO) categories enriched in in vivo quiescent NSCs (qNSCs) and activated NSCs (aNSCs). Each dot represents a BMP4-increased (red) or decreased (blue) protein associated with a given category. The outer ring indicates the $\log_2$ fold change (FC), the height of the inner bars reflects GO term significance ($-\log_{10}$ FDR), and color indicates Z-score. **E.** Left: Venn diagram showing overlap between BMP4-upregulated proteins and genes transcriptionally upregulated in qNSCs vs. aNSCs in our bulk RNA-seq dataset (Belenguer et al., 2021a). Center: heatmap showing the expression levels of BMP4-upregulated proteins in in vivo NSC populations. Right: $\text{Log}_2$-normalized mRNA expression levels of BMP4-upregulated proteins across qNSCs, pNSCs, and aNSCs. **F.** Bar plot of matrisome proteins upregulated in BMP4 vs. control samples, indicating $\text{Log}_2$Fold Changes (FC). Adjusted p-values are indicated. Bars are colored by matrisome category (Shao et al., 2022). **G.** Left: heatmap showing the expression of ECM-receptors from KEGG interaction pathway in BMP4 and control samples; significant adjusted p-values are indicated. Right: heatmap illustrating the in vivo expression of BMP4 upregulated receptors across NSC populations. Significant adjusted p-values between qNSC/aNSC are indicated in the heatmap. Scale bars: 10 $\mu$m.

### Supplementary Figure 3

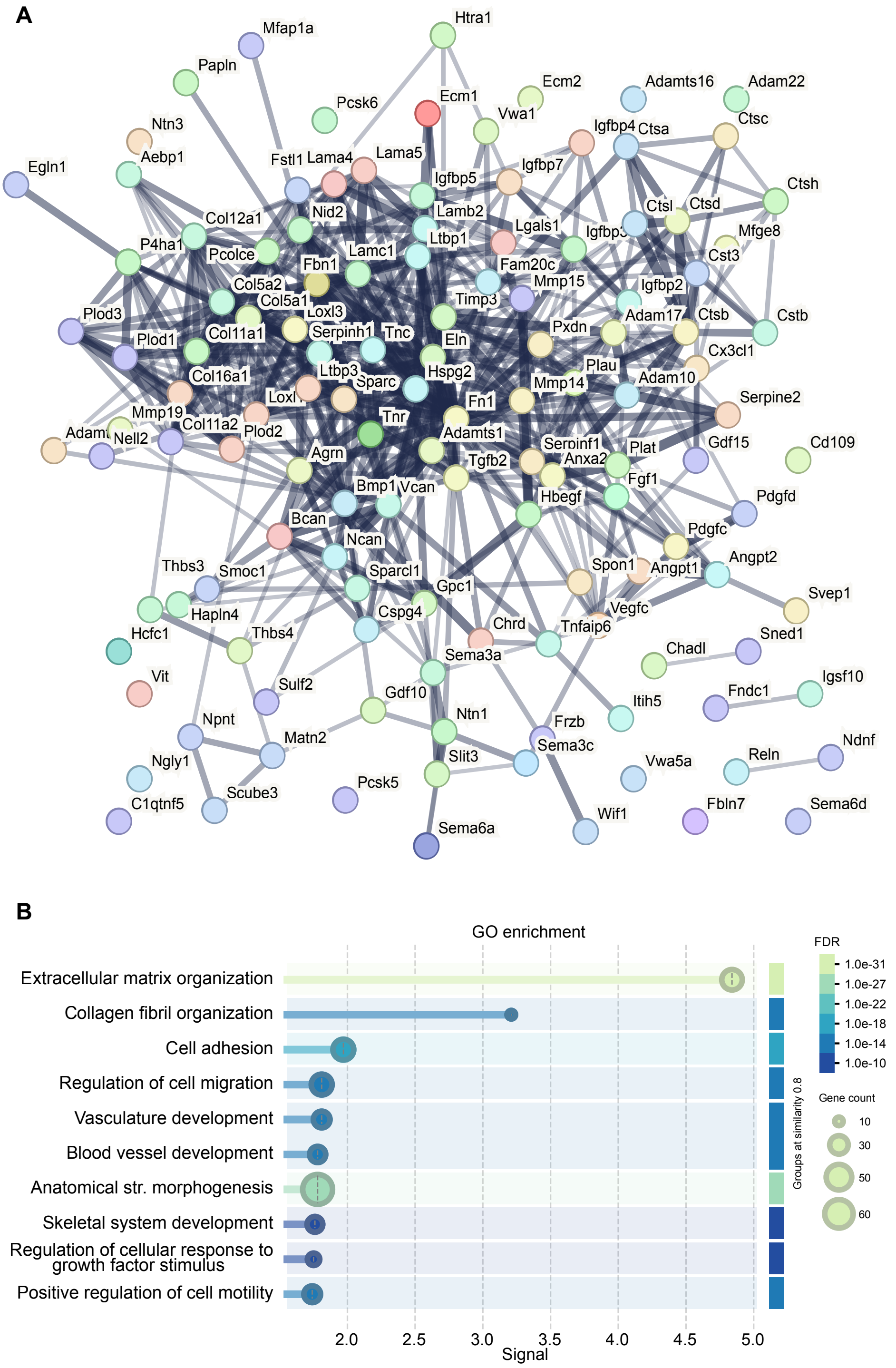


**Supplementary Figure 3. STRING analysis of deposited bMtx proteins. A.** STRING interaction network of matrisome proteins identified in bMtx. Nodes represent individual proteins, and edges represent protein–protein associations predicted or confirmed by experimental evidence. Line thickness indicates the strength and confidence of the interaction. **B.** Gene Ontology (GO) enrichment analysis of matrisome bMtx proteins. STRING-derived signal score reflecting enrichment strength is indicated. Circle size denotes the number of proteins associated with each GO term, and color intensity represents the statistical significance (FDR). GO terms are grouped by functional similarity (threshold 0.8) to reduce redundancy and highlight biologically coherent categories.

### Supplementary Figure 4

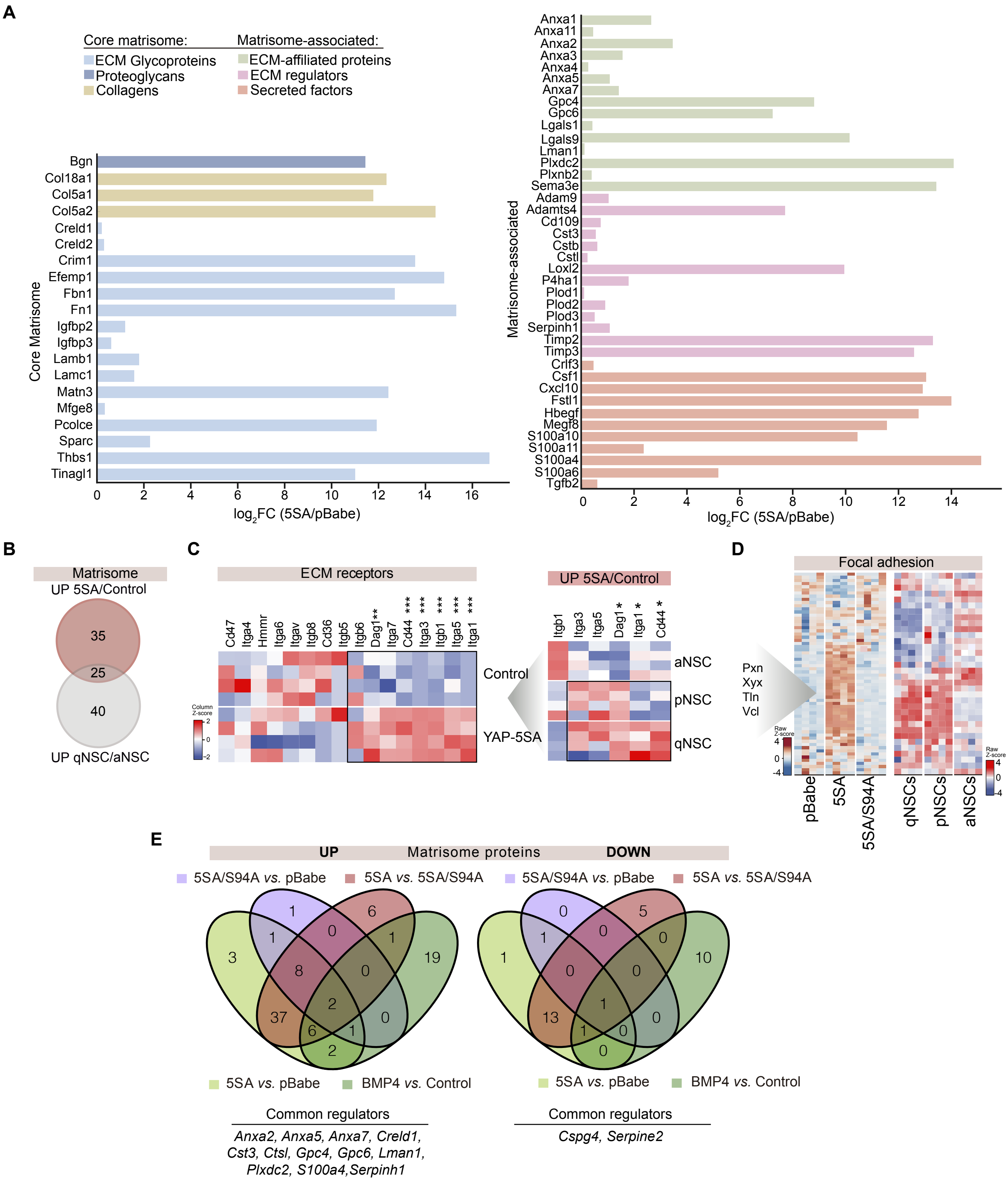


**Supplementary Figure 4. YAP activation promotes expression of matrisome and adhesion-related proteins. A.** Bar graph showing matrisome proteins significantly upregulated in YAP-5SA vs. control (pBabe). Proteins are categorized as core matrisome or matrisome-associated components. Bar length indicates $log_2$ fold change (FC), and colors represent specific matrisome categories. **B.** Venn diagrams showing the overlap between matrisome proteins upregulated in YAP5SA/pBabe and matrisome genes upregulated in qNSC/aNSC. **C.** Left: Heatmap displaying protein expression of ECM receptors from the KEGG ECM–receptor interaction pathway in YAP-5SA and control samples; asterisks indicate statistically significant changes. Right: Expression profile of significantly upregulated receptors in quiescent (qNSC), primed (pNSC), and activated (aNSC) NSC populations; asterisks indicate statistically significant changes. **D.** Left: Heatmap showing the expression of the focal adhesion gene signature (Nardone et al., 2017) across control, YAP-5SA, and YAP-5SA/S94A conditions. Right: Normalized expression of focal adhesion-associated proteins upregulated in YAP-5SA within NSC subpopulations in vivo. **E.** Venn diagrams depicting the overlap of upregulated and downregulated matrisome proteins across YAP constructs and the BMP4-induced proteome. Proteins commonly regulated by both YAP activation and BMP4 treatment are listed.

### Supplementary Figure 5

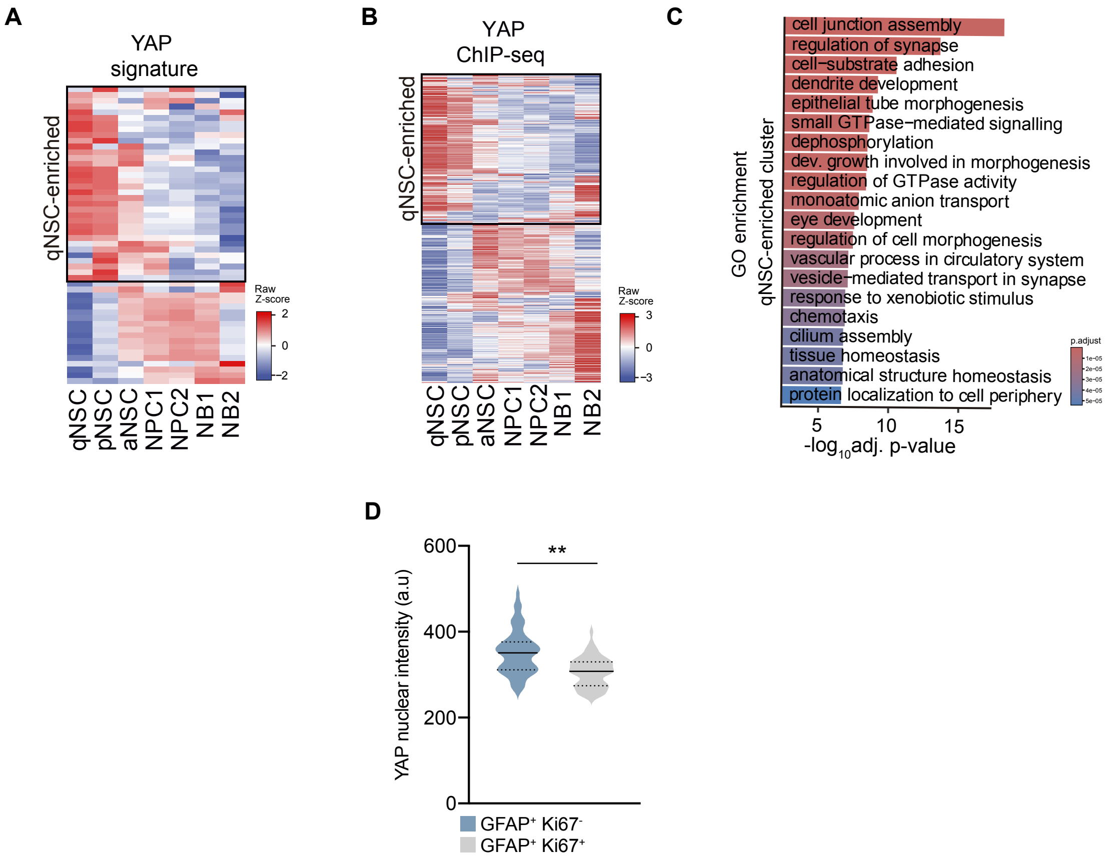


**Supplementary Figure 5. Enrichment of YAP transcriptional targets in quiescent NSCs. A.** Expression profile of the YAP transcriptional signature (Cordenonsi et al., 2011) across neurogenic lineage populations. Genes enriched in the qNSC cluster are highlighted. **B.** Chromatin immunoprecipitation sequencing (ChIP-seq) analysis of YAP binding sites (Nardone et al., 2017) evaluated in neurogenic lineage populations, showing a cluster of genes enriched in quiescent NSCs. **C.** Gene Ontology (GO) enrichment analysis (Biological Process category) of YAP ChIP-seq targets enriched in qNSCs. Statistical significance is indicated by adjusted p-values and enrichment scores. **D.** Quantification of YAP nuclear mean fluorescence intensity in vivo, measured in whole mount en-face preparations in GFAP$^{+}$ Ki67$^{+}$ and GFAP$^{+}$ Ki67$^{-}$ cells.

### Supplementary Figure 6

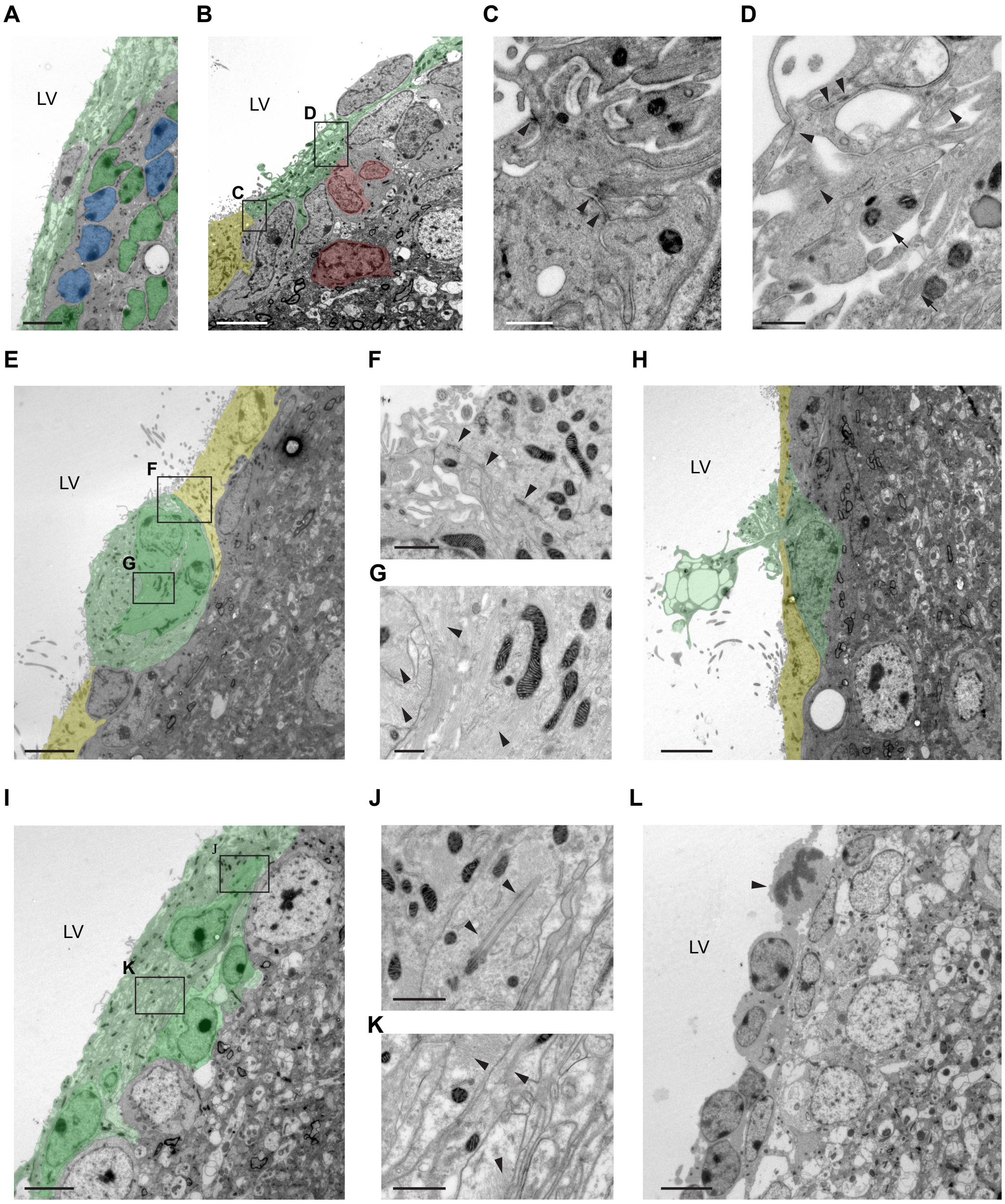


**Supplementary Figure 6. YAP/TAZ maintain the niche cellular organization and NSC polarity. A.** Increased cellularity in cYAP$^{KO}$/TAZ$^{HET}$ mice. B cells are denoted in green, progenitors in blue and astrocyte projections covering the ventricle are indicated in lighter green. **B.** V-SVZ panoramic view showing astrocytes (green) in contact with an ependymal cell (yellow) lining the ventricle. Proliferating cells are false-colored in red. Scale bar: 5 μm. **C.** Detailed view of the contact of the ependymal cell with the B cell. Arrowheads indicate junctions between the two cell types. **D.** Magnification of the astrocytic entangled projection contacting the ventricle. Arrowheads point to junctions between astrocytic projections and arrows indicate glial filaments. **E.** Panoramic view of two B cells in contact with the ventricle, disoriented and surrounded by astrocyte projections (green) immersed between ependymal cells (yellow). **F.** Detailed view of the contact of the ependymal cell with the B cell. Arrowheads indicate junctions between the two cell types. **G.** Arrowheads point to abundant glial filaments oriented in different directions. **H.** Panoramic view showing a large elongated B cell protrusion into the ventricle emanating from the V-SVZ (green) between the ependymal cells lining the ventricle (yellow). **I.** B cells covering the ventricle (green). **J.** Detailed view of missoriented astrocytic projection displaying a primary cilium (delimited by arrowheads) towards the V-SVZ tissue instead of contacting the ventricle. **K.** Interwoven and overlapping astrocytic projections containing abundant glial filaments oriented in different directions indicated by arrowheads. **L.** Loss of V-SVZ structure, B cells and progenitor in contact with the ventricle. Arrow points to a dividing cell in metaphase. LV: lateral ventricle. Scale bars: 5 μm (A,B,E,H,I); 500 nm (C,D,G); 1 μm (F, J, K).
